# Structural variation in repeat elements is widespread in normal human tissues and in tumorigenesis

**DOI:** 10.64898/2026.09.22.753313

**Authors:** Akshaya V. Annapragada, James R. White, Hope Orjuela, Adrianna Bartolomucci, Alice C. Eastman, Shashikant Koul, Kaui P. Lebarbenchon, Daniel C. Bruhm, Sarah Short, Keerti Boyapati, Noushin Niknafs, Carter Norton, Vishruth Girish, Nicholas A. Vulpescu, Sofia Velculescu, Julia Velculescu, Vilmos Adleff, Andrew Nelson, Zachariah H. Foda, Boris Winterhoff, Ronny Drapkin, Michael C. Schatz, Jillian Phallen, Robert B. Scharpf, Victor E. Velculescu

## Abstract

Somatic mosaicism contributes to genomic variation, yet postzygotic structural variants remain under-characterized. We performed long- and short-read WGS from multiple individuals (n=47 normal tissues; n=168 samples) and identified mosaic structural variants in all individuals and germ layers, impacting a median 285.2 kb/genome. Nearly half of breakpoints were independently validated, with tissue distributions reflecting both early and late developmental origins. Most mosaic variants were repeat-mediated and 8.3% overlapped functional elements, an enrichment compared to germline variants. To extend these analyses in samples where long-read sequencing is infeasible, we measured repeat alterations from short-read sequencing, recapitulating mosaic tissue-specific differences. We characterized tumor- and tissue-specific variation in repeats across 15 cancer types and found tumor-related repeat variation to be similar in scale to that of normal mosaic variation. Tracking repeat changes in cell-free DNA provided a noninvasive approach for tumor monitoring. Our analyses revealed widespread repeat-driven structural variation in health and disease.

**HIGHLIGHTS:**

- Multi-tissue long-read sequencing reveals widespread mosaic structural variants in normal tissues.
- Mosaic structural variation is largely repeat-mediated and has functional consequences.
- Alignment-free approaches enable analyses of repeat elements and large structural variants from short-read sequencing.
- Structural variation in repeat elements further occurs throughout tumorigenesis and may be detected in tumor tissues as well as through cell-free DNA liquid biopsies.

## INTRODUCTION

Genetic mosaicism resulting from somatic mutations in subsets of cells in the human body have been documented in normal aging, environmental exposures, and disease processes, including carcinogenesis, autoimmunity and neurodegeneration^1–6^. Historically, somatic sequence changes were thought to be largely restricted to clonal expansions within a differentiated tissue and serve as the predominant drivers of genomic variation outside the germline. However, postzygotic mutations may also begin in the earliest cell divisions after fertilization^7,8^, leading to somatic mosaicism across tissues^1,5,9^ and developmental stages^10,11^. The very earliest mutations lead to subclonal variants in tissues from all germ layers, while mutations later in development or adult life exhibit patterns of tissue-specific mosaicism restricted to single tissues or groups of embryologically related tissues^12^. Additional mosaic variants arising in a single cell of a differentiated tissue and then clonally expanding may lead to tumorigenesis^13–15^.

Most studies of genetic mosaicism have focused on single nucleotide variants (SNVs), small insertions and deletions (indels, <50 base pairs) or large-scale copy number alterations (CNA), using these to reconstruct cellular phylogenies from zygote to adult^8,11^, characterize individual organs^16–19^, or identify early signatures of cancer^3,20^. Larger structural variants (SVs, ≥50 base pairs) are of growing interest^10,21^, but are only beginning to be studied in multi-tissue samples from the same donor^9^, and their occurrence and mechanistic origins remain incompletely understood^10^. SVs intrinsically impact large genomic segments and may drive more variation than SNVs and indels, yet remain poorly characterized due to challenges resolving their structure and localizing breakpoints in conventional short-read sequencing^21–25^. In mosaic variants with subclonal variant allele fractions (VAFs), often below 5%, the scarcity of variant reads further limits detection^21^.

Repetitive genomic regions are thought to represent hypermutable hotspots for SV formation^26,27^ via several potential mechanisms including LINE-1 mediated target-primed reverse transcription (TPRT)^28^, non-allelic homologous recombination (NAHR)^29^, microhomology-mediated repair and replication-based template switching (MMEJ/FoSTeS/MMBIR)^30^, and other related replication repair and recombination pathways^31^. However, repeats are among the most complex regions of the genome to resolve due to high background germline variation and low mappability^32,33^, and functional roles for these largely noncoding genomic elements are incompletely understood^32,34–36^. Long-read sequencing has enabled the resolution of complex variants within repetitive sequences, leading to detection of somatic SVs in tumor tissues with high VAFs^37–39^, discovery of putative mosaic SVs in single tissues^40–42^, and identification of *de novo* gonosomal mosaicism in trio studies^41,43^. However, high input DNA requirements of long-read sequencing^44^ have limited widespread use and multi-tissue studies. We previously developed an approach, ARTEMIS, to construct alignment-free repeat landscapes from conventional short-read sequencing with low DNA inputs^45^. This complementary approach allows for global quantification and classification of over 700 repeat element types whose changing levels in the repeat landscape correlate with genomic instability and the presence of SVs including LINE-1 mediated deletions, gene amplifications and complex breakpoints. ARTEMIS also allows for analysis of repeat landscapes in highly fragmented cell-free DNA (cfDNA), an analyte of interest for noninvasive liquid biopsies in cancer detection and monitoring^46^.

In this study, we present a comprehensive characterization of mosaic structural variation through long- and short-read sequencing of multiple tissues with distinct embryological origins from the same individuals (Figure 1A). We examined repeat elements as a source this mosaicism, as well as the functional consequences of mosaic structural variants. We then expanded these analyses to compare normal tissue-specific repeat mosaicism with tumor-related changes in repeat elements, and examined tumor evolution from ovarian pre-cancer lesions to tumors, both in tissues and through liquid biopsies. We hypothesized that these analyses would reveal previously underappreciated SV mosaicism and mechanisms of genome-wide tissue-specific changes across development and tumorigenesis.

**Figure 1.**
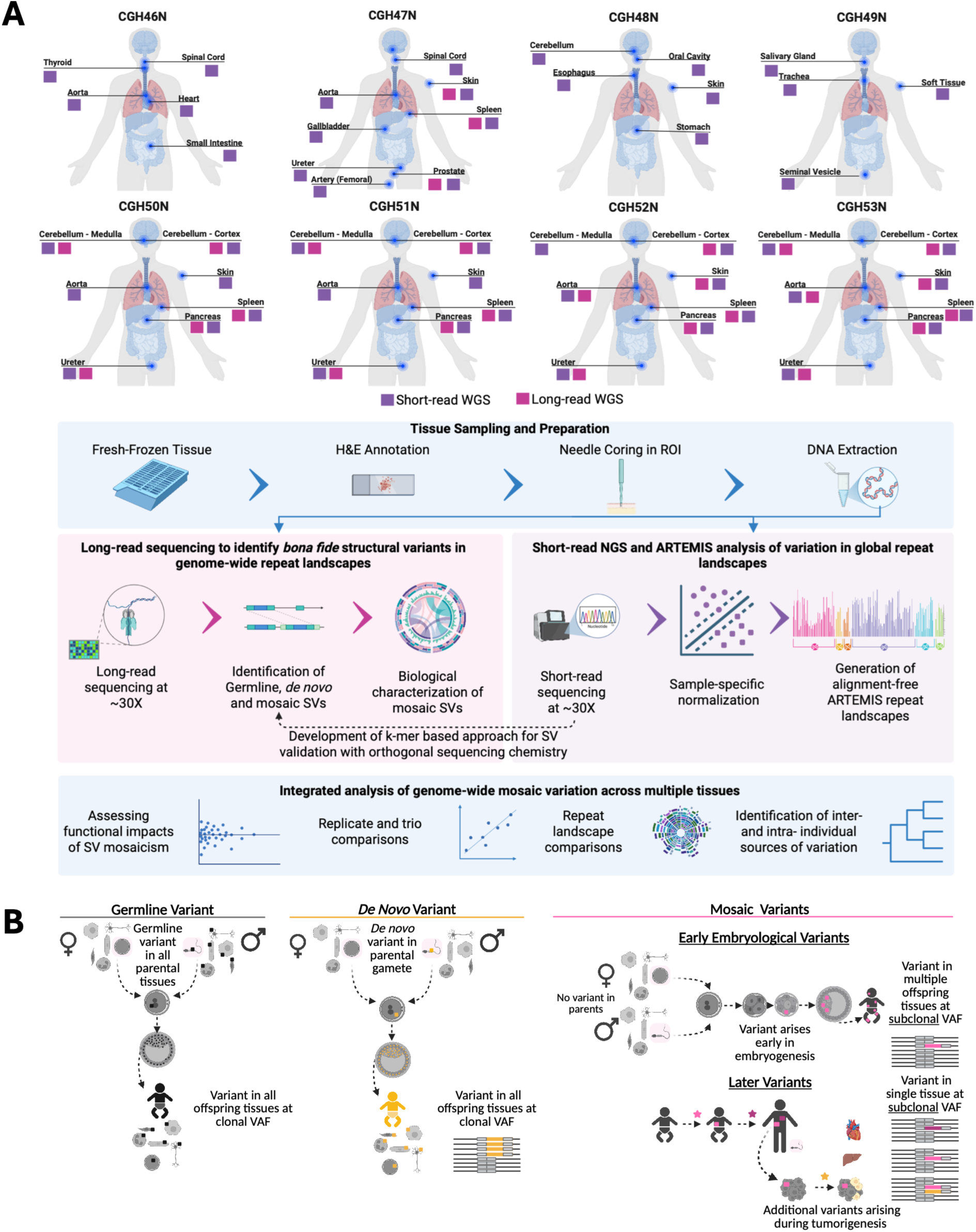
Study design for investigation of structural variants and mosaic repeat landscape variation across multi-organ samples. **(A)** Tissue samples were obtained from multiple organs of eight individuals, and both short and long read whole genome sequencing were performed. This data was used to characterize *bona fide* structural variants and changes to global repeat element landscapes which may be driven by previously underappreciated mosaicism. **(B)** Schematic of mechanisms behind occurrence of germline, *de novo*, and mosaic variants. Germline variants are passed from parent to offspring, while *de novo* variants are somatic variants arising during life in parental gametes that are then passed to children at clonal VAF. Mosaic variants are somatic variants occurring in offspring. For early embryological mosaic variants, tissue distribution and VAF are determined by timing of variant occurrence with respect to gastrulation, though VAF is generally expected to be subclonal as the variant initially occurs only in a single cell post-fertilization. Clones arising pre-gastrulation may appear in most or all tissues across germ layers, while post-gastrulation variants may occur only in tissues of a specific lineage. Later variants occurring after birth are typically restricted to a single tissue and appear subclonal on bulk sequencing due to spatial localization within the tissue. Subsequent disease-associated variants arising in tumorigenesis are constrained to a single tissue. See also Figure S1.

## RESULTS

### Identification of mosaic, de novo and germline structural variants in multiple tissues using long-read sequencing

Mosaic SVs are postzygotic somatic events and can arise early in embryogenesis across multiple tissues or later in life in one or a few tissues, typically at subclonal VAFs (Figure 1B). To examine somatic mosaic variants throughout development and adult life, we obtained 47 fresh-frozen tissue blocks from eight individuals (JHU cohort, range 4 - 8 tissue types per individual) encompassing 21 organs, including aorta (n=6 individuals), cerebellum (n=5), skin (n=6), spleen (n=5), pancreas (n=4) and ureter (n=5) (Figure 1A, Tables S1 and S2). The analyzed individuals were aged 37 - 73 years and died of non-cancerous causes. For most individuals, we prepared two biological sample replicates of each tissue by macro-dissecting distinct hematoxylin and eosin (H&E) annotated regions, extracting DNA separately, and processing these as separate samples (Figure S1).

For samples with sufficient extracted DNA, we created genomic libraries and sequenced these using nanopore long-read sequencing with a median 31.5X coverage and 6.4 kb N50 (n=47 samples, representing 23 tissues from 5 individuals) (Table S3). We aligned reads to the gapless reference genome (T2T-chm13)^47^, performed haplotype-aware phasing, and identified SVs using a breakpoint graph-based algorithm^37^. After filtering low confidence calls or likely artifacts, we retained 48314 SVs >=50bp (Tables S4, S5, S6, S7, and S8). Most SVs (n=43745) were common germline events also found in the 1000 Genomes data^48^. Of germline SVs not occurring in 1000 Genomes or in multiple unrelated individuals, variants with near-clonal VAFs (> 0.25) in any tissue sample were classified as rare or *de novo* (n=1959) germline variants. Variants found only at subclonal VAF (≤0.25, a conservative threshold selected to minimize false positive calls from germline and *de novo* changes^41^) were designated as putative mosaic variants (n=2610) (Figures 1B and 2A, Materials and Methods). Mosaic SVs were found at subclonal VAFs as low as 0.02 (median 0.07, IQR 0.06-0.09), while the distributions of germline and de novo SV VAFs clustered around 0.5 and 1.0 as expected (Figure 2B).

**Figure 2.**
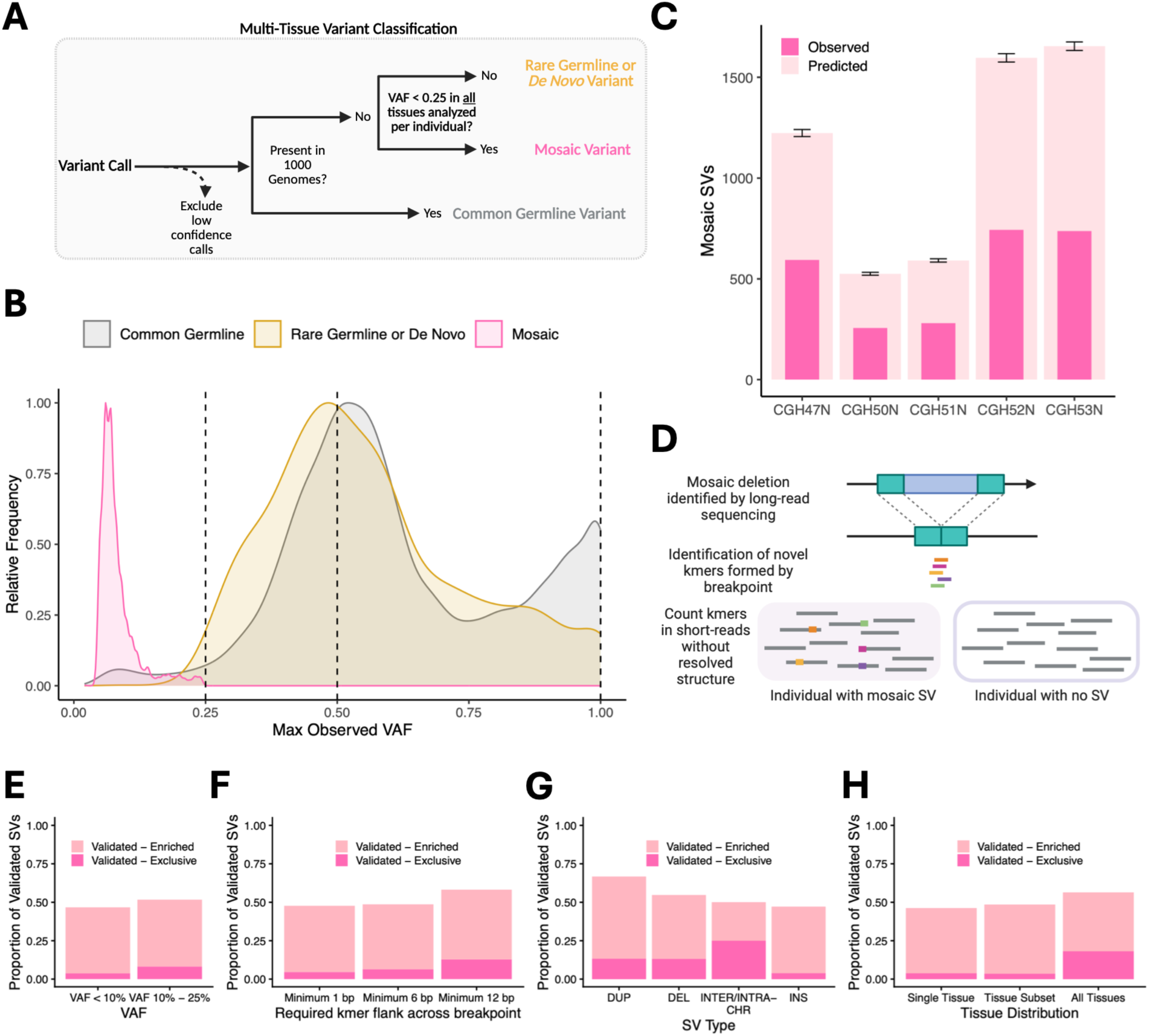
Discovery and validation of mosaic SVs. **(A)** Algorithm for multi-tissue variant classification. High quality variants also found in the 1000 Genomes call set are classified as common germline events. Among other variants, those found at a clonal VAF (>0.25) in any tissue sample are classified as Rare Germline or *De Novo* variants. Variants found at subclonal VAF (≤0.25) in all tissue samples in which the variant is called are classified as mosaic variants. **(B)** Distribution of observed VAFs across structural variants called in the JHU Cohort. As expected, Germline and De Novo SVs show VAF peaks near 0.5 (heterozygous) and 1.0 (homozygous), while mosaic VAFs are evident across a subclonal range from 0.02 - 0.25). **(C)** Observed and predicted numbers of true mosaic SVs for each individual demonstrate that the true number of mosaic SVs is likely higher than the observed number, due to sensitivity limits imposed by moderate sequencing coverage. Estimates are derived from binomial simulations of variant calling informed by precision and recall estimates at 30X coverage computed using the SMaHT-MIMS benchmark. Error bars show +/− 1 standard deviation on the range of predicted SV numbers. **(D)** Kmer-based short-read sequencing approach for orthogonal validation of mosaic SVs discovered with long-read sequencing. First, novel 24bp sequence (kmers) formed by SV breakpoints and not found within the chm13 reference genome are identified. These kmers are then counted in short-read sequencing reads from samples of the individual with the SV and unrelated individuals, and novel kmer abundance is compared. **(E)** Proportion of evaluable mosaic SVs validated using the kmer-based approach in (D), stratified by VAF. An SV is considered evaluable if atleast 1 novel breakpoint kmer is found in atleast 1 sample. An evaluable SV is considered Validated-Enriched if the median kmer count for that SV is higher in the individual with the variant than unrelated individuals. An evaluable SV is considered Validated-Exclusive if the median kmer count in the unrelated individuals is 0. **(F)** Proportion of validated mosaic SVs depending on the minimum required flank for novel kmers on either side of the SV breakpoint. A novel kmer is one not found in the chm13 reference genome and formed by a SV breakpoint. At a minimum 1bp, any novel kmer spanning the breakpoint is counted. For a minimum 6bp flank, atleast 6bp must be found on either side of the breakpoint. For a minimum 12bp flank, 24bp kmers must be centered across the breakpoint. **(G)** Similar to (E), but for different types of mosaic SVs. **(H)** Similar to (F), but for different tissue distributions of mosaic SVs. See also Figures S2, S3, S4, S5, S6, S7, S8, S9, and S10.

### Orthogonal validation of mosaic structural variants

To evaluate our SV classification approach in well-annotated reference samples, we called mosaic SVs from blood cell DNA of seven Genome in a Bottle (GIAB) individuals^49^ analyzed with long-read sequencing, and also identified SNVs and indels (<50bp). SVs were less common than SNVs and indels but impacted larger genomic segments (median length of 1 bp vs. 5 bp vs. 509 bp for mosaic SNVs, small indels, and SVs respectively), highlighting their biological significance for genome structure (Figure S2A). We then tested our classification approach using the SMaHT-MIMS mosaicism benchmark^21^, a synthetic mosaic DNA sample combining different individuals and sequenced to yield 49368 structural variants at a median VAF of 0.06 (IQR 0.01-0.46). For mosaic SVs called on this ONT long-read benchmark at 30X sequencing coverage, recall increased with VAF (4.7%, 25.3%, 43.6% and 54.5% for VAFs ≤1%, 1%-5%, 5%-10% and 10%-25% respectively), while precision remained uniformly high (range 87.2%-89.2% across the four VAF bins) (Figure S2B). Our approach misclassified only 3.6% of non-mosaic SVs as mosaic, and increased sequencing coverage yielded only modest improvements (Figures S2C and S2D). An analysis of potential technical and classification errors using two GIAB parent-offspring trios similarly revealed that rates of false positive mosaic calls were low, and likely driven by variant calling from a single tissue (blood) sample (Figures S2E and S2F). In the JHU cohort, mosaic variants could be confirmed across multiple samples and tissues, with median supporting read length of 6914 bp (IQR 3489 bp – 11988 bp) and median 66.7% of supporting reads assigned to the expected parental haplotype (Figure S3). Monte Carlo simulations of SV detection informed by the above benchmarking suggested that our observed 2,610 mosaic SVs across 5 individuals likely represented a conservative estimate of the extent of structural mosaicism, with the true number as high as 5590 mosaic SVs (range 525-1654 mosaic SVs per individual) (Figure 2C).

We aimed to experimentally validate the identified long-read mosaic SVs using standard next generation Illumina sequencing (150 bp paired-end sequencing). Because such short-read sequencing cannot reliably resolve large structural variants^25,26,37^ and changes in repeat-rich genomic regions^32^, we developed a kmer based approach for validating mosaic SVs (Figure 2D). We hypothesized that newly created short sequences (e.g. 24 bp) formed by the SV sequence breakpoints not found in the reference genome may be present in individuals with SVs but absent or present at lower frequencies in unrelated individuals. Using this approach, we considered an SV to be validated if the frequency of novel breakpoint kmers found in sequence data of tissue samples from the individual with the putative SV was greater than that observed in samples from unrelated individuals (Figure S4A). We found that 47.7% of mosaic SVs identified in long-read sequencing that could be evaluated were validated using this approach (Figure 2E). Increasing the flank sequences on either side of the SV breakpoint decreased the proportion of evaluable mosaic SVs (Figures S4B and S4C) but increased the successful validation rate, likely because such kmers were more likely to be truly unique to the SV (Figure 2F). Orthogonal SV validation rates remained above 46% for all types of mosaic SVs and regardless of tissue distribution within an individual (Figures 2G and 2H).

### Landscape of mosaic structural variants across human development

The identified 2610 mosaic SVs comprised large insertions (n=2071), deletions (n=398), duplications (n=39), and inter-(n=1) and intra-(n=101) chromosomal rearrangements (Figures 3A, S5, S6, S7, S8, S9 and S10). Complex chromosomal rearrangements were more common among mosaic SVs than germline or de novo SVs (3.9% of mosaic SVs vs. <1.02% for rare germline/de novo and common germline SVs, p<3.7×10^−9^, Chi-square test).

**Figure 3.**
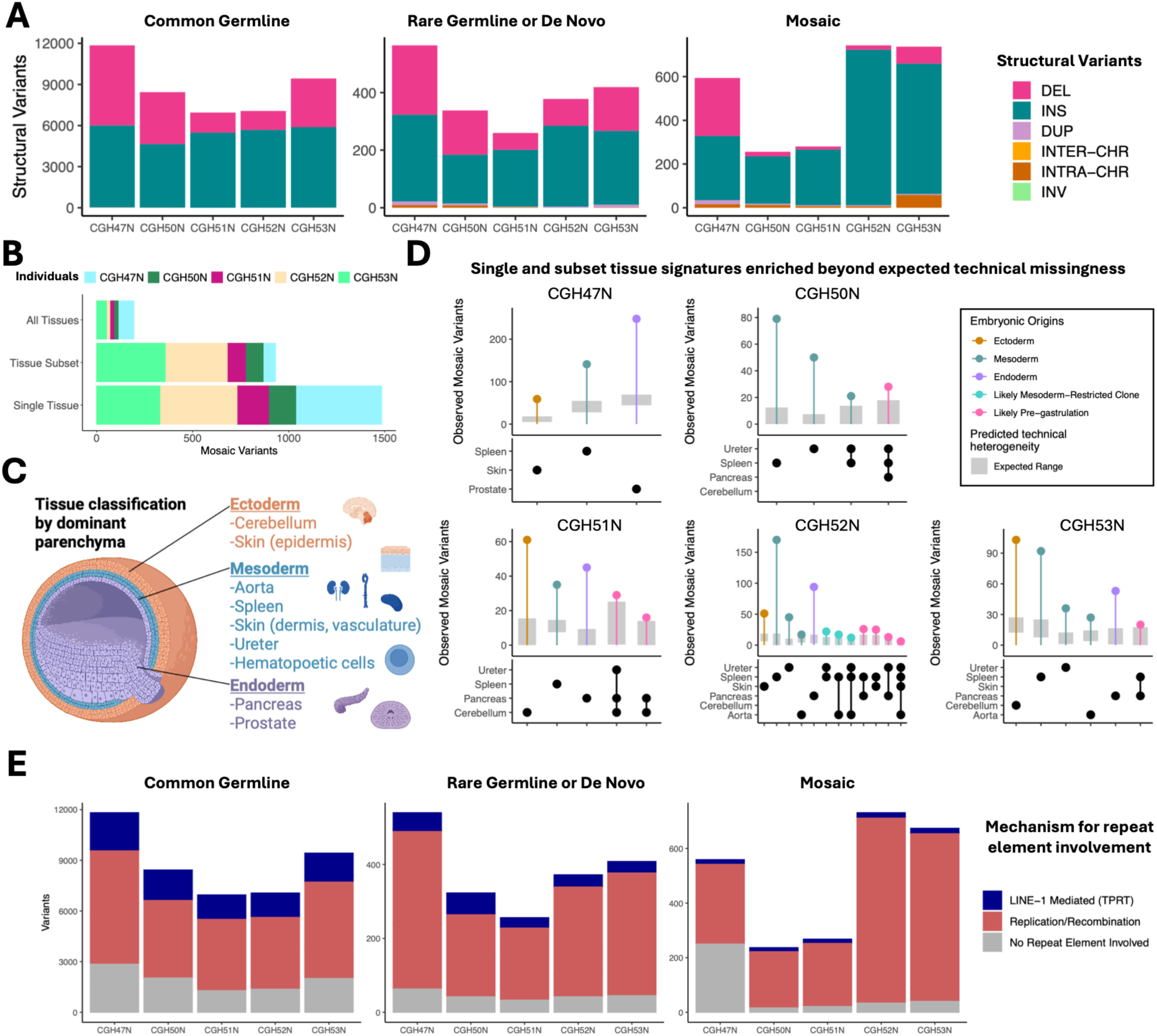
Genome-wide analyses across multiple matched tissues reveal that mosaic SVs occur across developmental lineages and exhibit characteristics of repeat element involvement. **(A)** SV types within the JHU Cohort for putative common germline, rare germline/*de novo* and mosaic variants. DEL, deletion. INS, insertion. DUP, duplication. INTER-CHR, inter-chromosomal rearrangement. INTRA-CHR, intra-chromosomal rearrangement. INV, inversion. **(B)** Tissue distribution for mosaic variants. The majority of mosaic SVs are found a single tissue at VAF ≤0.25. Some also occur at subclonal VAF in multiple or all tissues, indicating early embryonic origins. **(C)** Germ layer origins of tissues studied, classified by dominant parenchymal origin. For each individual, tissues originating from all three germ layers are represented. **(D)** Observed numbers of tissue-restricted mosaic SVs as compared to numbers predicted due to technical missingness alone. All statistically enriched tissue signatures are shown, colored according to their hypothesized biological origins (pre-gastrulation, lineage-restricted or single germ-layer/tissue). **(E)** Predicted mechanisms of occurrence for INS and DEL SV calls in the JHU cohort reveal that most mosaic insertions and deletions involve repeat elements. Repeat-involved mechanisms include LINE-1 mediated target-primed reverse transcription (TPRT) and processes involving replication and recombination including variable number tandem repeat (VNTR) insertions and deletions, non-allelic homologous recombination (NAHR), and microhomology-mediated repair and replication-based template switching (MMEJ/FoSTeS/MMBIR) See also Figure S11.

We approximated the developmental timing of mosaic SVs from their tissue distribution. Variants present across all tissues likely arose pre-gastrulation, while lineage-restricted variants would be expected to occur during post-gastrulation embryogenesis, and single-tissue variants would likely occur in later development or postnatal events. In the JHU mosaic SV call set, we observed that 7.4% of subclonal SVs were in all sampled tissues within an individual, 35.7% were in subsets of tissues, and 56.9% were in a single tissue (Figure 3B, Tables S4, S5, S6, S7, and S8), indicating that while most large mosaic SVs occur later in life and impact single tissues, a substantial fraction may originate during embryonic development.

To identify tissue-specific mosaic changes, we examined read-level allele counts for each variant using a model that considered tissue-specific coverage and technical heterogeneity in sequencing and variant calls^23,49^ (Figures S11A and S11B). From these analyses, we observed biologically plausible tissue- or lineage-specific restriction, including clones confined to mesodermal tissues or spanning multiple germ layers, consistent with post-gastrulation and early embryonic origins, respectively (Figure 3C and 3D). Among the tissues analyzed, we observed that spleen-only mosaic variant calls were enriched in all individuals, perhaps due to hematopoietic mosaicism in infiltrating blood cells^50,51^. Cell-type-of-origin analyses of long-read sequence data using 5-methylcytosine (5-mC) and 5-hydroxymethylcytosine (5-hmC) modifications confirmed that spleen tissue samples contained a median 62.1% infiltrating blood cells, while other tissues were largely composed of tissue-specific cell types (Figure S11C). Cerebellum-only mosaic variants were enriched in 2 of 4 individuals with this tissue sequenced, consistent with prior reports of brain specific mosaicism^52–54^. Skin-specific mosaic SVs were enriched in 2 of 3 individuals with this tissue sequenced, suggesting that reports of high single nucleotide mutational burden in skin^1,55^ may also apply to structural variants. We further observed an enrichment of single tissue SVs in other tissues not previously reported to exhibit high mutational burdens^1,56^ or structural variant mosaicism^9^ including pancreas (3 of 4 individuals sequenced), suggesting that SVs may accumulate in normal tissues throughout life at previously underappreciated rates.

### Enrichment of mosaic structural variants within genomic repeat elements

To understand mechanisms underlying structural mosaicism, we annotated insertions and deletions in the JHU call set for signatures of repeat-element–mediated rearrangements^28–31,48^, including LINE-1–mediated events via target-primed reverse transcription (TPRT) and replication/recombination mechanisms (e.g., NAHR, FoSTeS/MMBIR). Most SVs carried one or more of these signatures (78.2%, 88.2%, and 85.2% of common germline, rare germline/de novo, and mosaic SVs, respectively) (Figure 3E, Tables S9 and S10). Given that approximately 50% of the genome contains repetitive sequences^32^, this represents significant enrichment across all three groups (p<2.2×10^−16^, one-sided binomial test), consistent with repeat-rich regions being hotspots for postzygotic SVs due to homology-driven replication errors and mobile element activity.

We hypothesized that SVs may be enriched in in repetitive regions of the genome due to lower density of essential coding sequence^32,34,35^. To evaluate this selective constraint hypothesis, we analyzed the genomic sequences of 89,432 19 bp simple guide RNAs (sgRNAs) used to introduce double stranded breaks (DSBs) in genome-wide CRISPR essentiality screens across 318 cell lines conducted by the DepMap consortium^57^. Across cell lines and gene targets, sgRNA-induced deleterious DSBs were attenuated when the sgRNA cut-sites occurred within annotated repetitive elements (Figures S12A and S12B), suggesting that repeats may serve as a “buffer” during early development, allowing cells to survive mutagenic events that could otherwise be lethal. The mosaic SVs found in the JHU Cohort formed kmers at their breakpoints with homology to these annotated repeat elements, with each mosaic SV forming kmers mapping to a median of 110 repeat element types (IQR 8-728) from the LINE, SINE, LTR, Satellite and Transposable element families (Figures S12C and S12D), suggesting that repeat mediated attenuation of deleterious DSBs may promote persistence of mosaic SVs.

### Mechanisms for repeat element involvement in mosaic structural variants

LINE-1 elements remain among the only active retrotransposons capable of mobilizing SINEs and LINEs via TPRT^28^, and they accounted for 3.6% of mosaic SVs in the JHU call set (Figures 4A and S13A). Other repeat-involved mosaic SVs reflected variable number tandem repeats (VNTRs) or transposable element duplications (72% and 9.6%, respectively), consistent with NAHR or template-switching (FoSTeS/MMBIR) mechanisms. As an example, we observed a canonical L1-mediated SINE AluY insertion at chr9:78,573,892 with hallmark features including target-site duplication and insertion of a 270 bp Alu-like element upstream of a poly-A tail (Figure 4B). This mosaic SV was confirmed by two variant callers and was present at subclonal VAF (1.3–6.9%) across all studied tissues, including prostate, skin, and spleen, in both biological and library replicates (Figures 4C and S5A). As a second example, we identified a mosaic interspersed duplication at chr5:178,593,183 of a 2.53 kb AluSx/MIR block copied from chr5:176,649,942, with short templated-sequence insertions (16 bp left; 5 bp right), minimal microhomology (1 bp left; blunt right), and no target-site duplication (Figure 4D), consistent with a replication-based template switching mechanism (FoSTeS/MMBIR). Uniform coverage at donor and acceptor sites further supports a copy–paste event observed at subclonal VAF in pancreas, spleen, and ureter but absent from cerebellum, suggesting a mesoderm/endoderm-restricted mosaic variant that likely arose near gastrulation, shortly after ingression of the neural crest^11^ (Figures 4E and S5B). These mosaic developmental origins to our knowledge have not been previously documented for large SVs.

**Figure 4.**
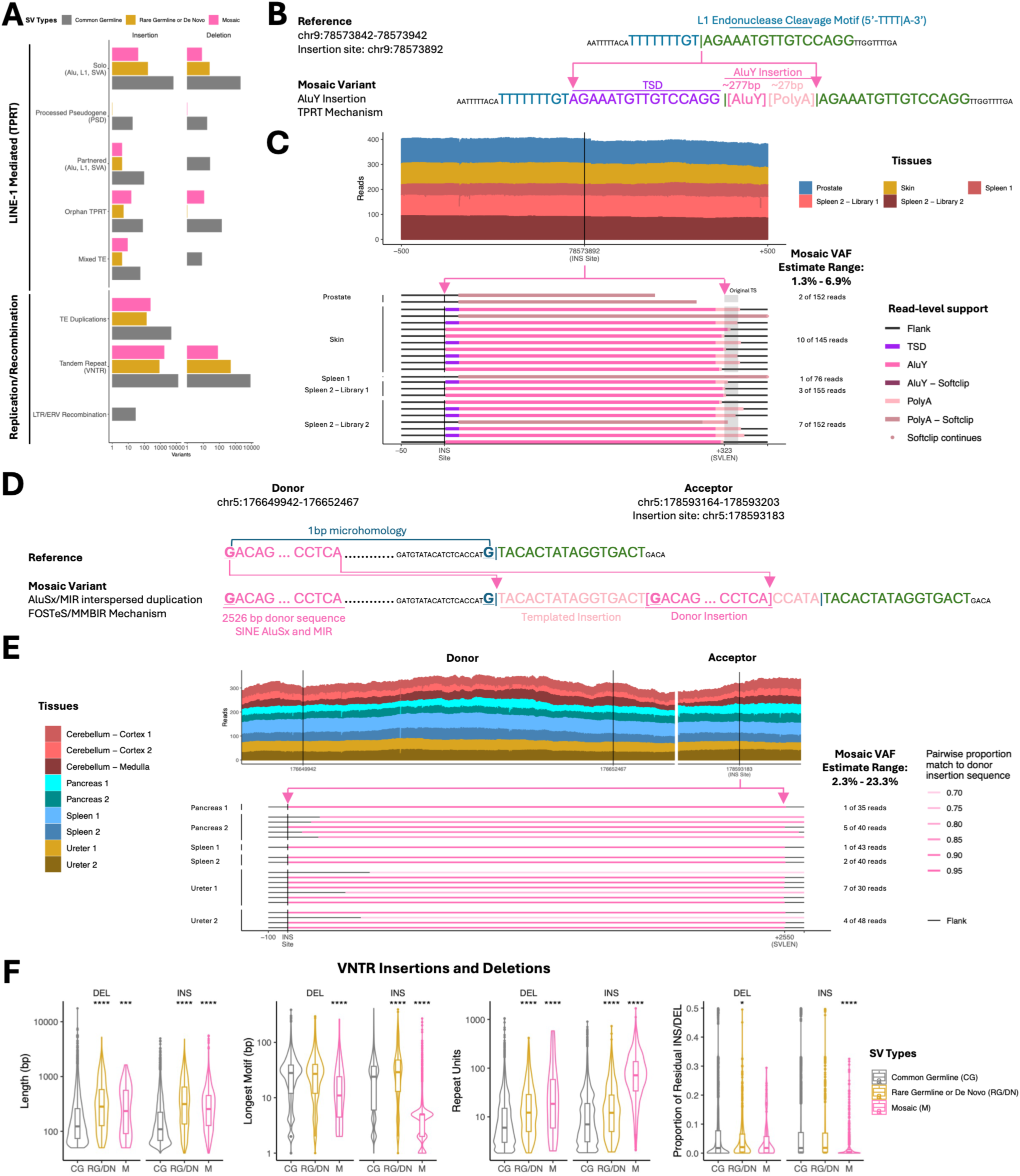
Repeat element mediated mechanisms drive mosaic SVs across tissues. **(A)** Detailed annotation of mechanism for repeat-element involved insertion and deletion SV calls in the JHU Cohort. Signatures of both LINE-1 mediated and replication/recombination mechanisms are present across germline, *de novo* and mosaic SVs. **(B)** Example of a mosaic AluY insertion in the JHU Cohort with characteristics of LINE-1 mediated target-primed reverse transcription (TPRT), including target site duplication (TSD), L1 endonuclease cleavage motif, and an inserted AluY element with poly-A tail. **(C)** Read-level support for this mosaic LINE-1 mediated AluY insertion demonstrates uniform coverage across all tissues with long-reads spanning the breakpoint and target site (TS) and containing target site duplication (TSD), poly-A tail and AluY homology. The SV is observed at subclonal VAF in all studied tissues of this individual including prostate, skin and spleen with library and biological replicates. **(D)** Example of a mosaic interspersed duplication via a “copy-paste” mechanism in the JHU Cohort with characteristics of a replication-based template switching mechanism (FoSTeS/MMBIR), including templated insertion of an AluSx/MIR element and minimal microhomology. **(E)** Read-level support for this mosaic interspersed duplication demonstrates uniform coverage across all tissues at both the donor and acceptor site, with long-reads matching the donor sequence and spanning the acceptor breakpoint. The SV is observed at subclonal VAF in pancreas, spleen, and ureter but absent from cerebellum suggesting a lineage-restricted mosaic variant arising near gastrulation but shortly after neural crest ingression. **(F)** Characteristics of variable number tandem repeat (VNTR) insertions and deletions in the JHU Cohort. The total SV length (panel 1, leftmost), longest dominant motif (panel 2), number of repeat units (panel 3) and proportion of residual incomplete sequence (panel 4, rightmost) varies for common germline, rare germline/*de novo* and mosaic SVs, with the longest lengths and highest number of repeat units generally found in mosaic and rare germline/*de novo* SVs. See also Figures S12 and S13.

Overall, mosaic SVs were substantially longer than common germline SVs (median 202 bp vs. 144 bp, p<2.2×10^−16^), with the difference most pronounced among L1-mediated events (median 226 bp vs. 107 bp, p<2.2×10^−16^). Among SVs with L1-mediated TPRT signatures, size distributions showed peaks roughly at 300 bp and 6000 bp, corresponding to SINE and LINE element lengths (Figure S13B). For VNTR insertions, mosaic SVs had greater lengths than common germline SVs (median length 255 bp vs. 109 bp, p <2.2×10^−16^) and contained more repeat units (median 71.4 vs. 7.02 repeat units, p <2.2×10^−16^) though mosaic SVs had shorter dominant motif lengths and lower proportions of residual sequence (Figures 4F and S13C). For VNTR deletions, trends were similar, with mosaic and rare germline/de novo deletions affecting larger sequences than common germline variants (median 234 bp vs. 123 bp, p =0.0003).

### Genome-wide functional impact of mosaic structural variants

We next investigated the genome-wide distribution of mosaic SVs and their potential functional impacts. Early embryonic mosaic events found across tissues as well as germline and *de novo* SVs were significantly enriched in early-replicating GC-rich regions of the genome^58^ (p<7.2×10^−35^ for common germline, rare germline/de novo and mosaic SVs found in all tissues, Benjamini-hochberg corrected p-value from chi-squared goodness-of-fit test against a uniform distribution) (Figure 5A). These observations are consistent with the notion that genomic features of open chromatin and active transcription foster germline variability^59,60^ and may harbor SV mutational hotspots early in development. Germline, de novo and early developmental mosaic events were also enriched near telomeres, likely due to error-prone replication in regions enriched for tandem repeats and segmental duplications^61^ (p<2.19×10^−14^) (Figure 5A). In contrast, lineage- and tissue-restricted mosaic SVs arising later in development were uniformly distributed across early and late replicating regions and exhibited higher AT content, suggesting that as cells differentiate, somatic events may favor heterochromatic regions of the genome (p<1.37×10^−3^ for mosaic SVs found in tissue subsets or single tissues) (Figure 5A). All SVs were enriched in regions close to gene bodies (median 5.3 kb, range 0 – 23.5 Kb, p<4.2×10^−3^), consistent with mechanisms of recombination that favor gene- and GC-rich regions of the genome^62,63^.

**Figure 5.**
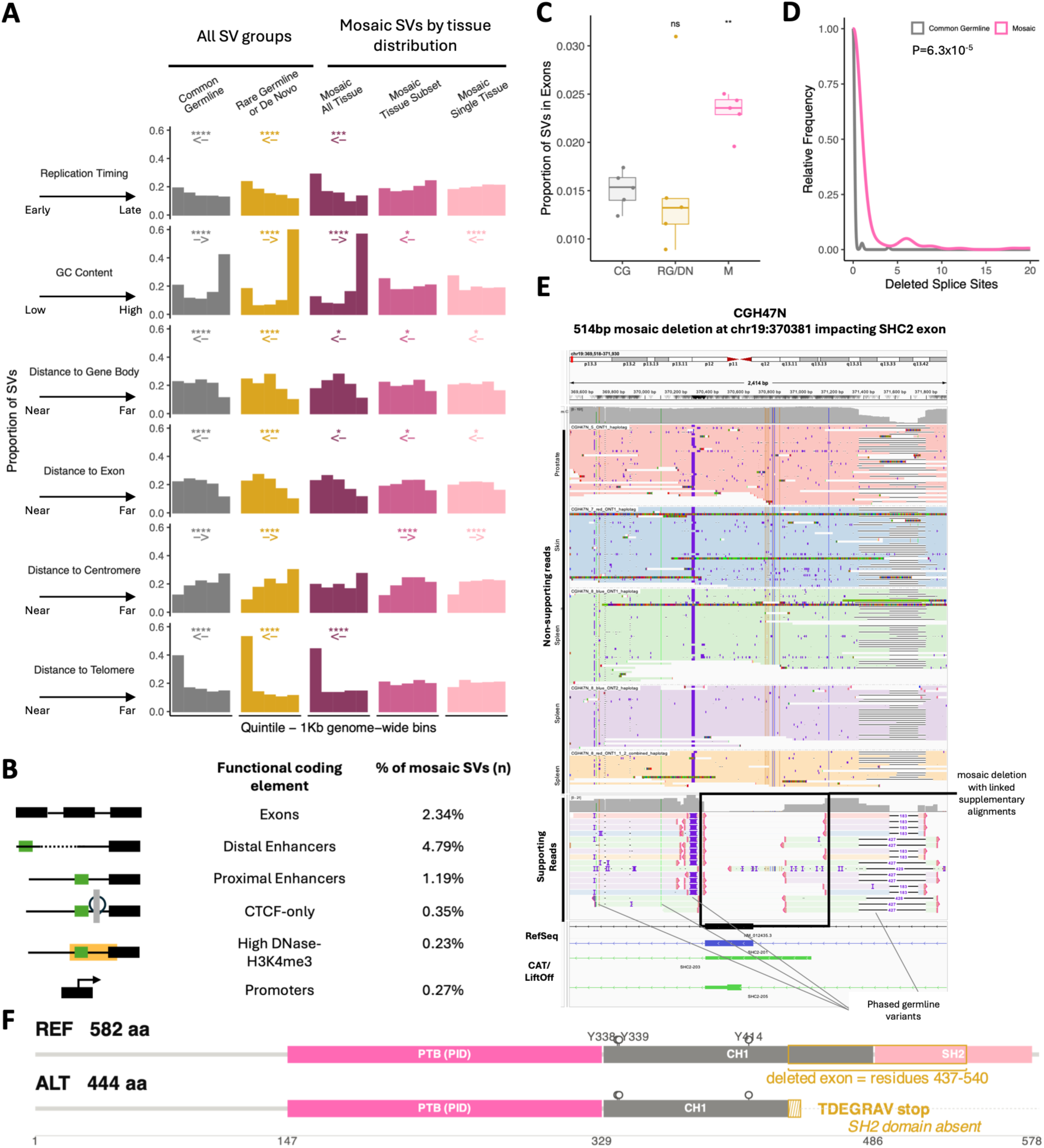
Genome-wide characteristics of SV breakpoints illustrate functional impacts of mosaic SVs. **(A)** The proportion of SVs overlapping 1Kb genomic bins stratified by quintile with respect to various genomic properties reveals non-uniform distribution of SVs across the genome. The Benjamini-Hochberg corrected p-value from a chi-squared goodness-of-fit test against a uniform distribution is computed for each variant type and genomic category (*, p<.05; **,p<.001;***,p<.0001;****,p<.00001) and the arrow shows the direction of distribution shift. **(B)** The percentage of mosaic SVs overlapping various coding elements in the genome highlights potential functional impacts. **(C)** The proportion of each group of SVs with breakpoints within exons reveals an enrichment of mosaic SVs disrupting coding exons as compared to common germline SVs. Each point represents one individual in the JHU Cohort. **(D)** Distribution of the predicted number of intron-exon splice sites lost due to mosaic SVs and a length and VNTR matched set of common germline SVs in the JHU Cohort. The rightward shift in the distribution for mosaic SVs shows that mosaic deletions are predicted to interfere with a larger number of splice sites than comparable germline deletions. **(E)** IGV screenshot of a mosaic deletion impacting an SHC2 exon. Non-supporting reads spanning the breakpoint are shown in a collapsed view, grouped and colored by tissue sample of origin. The reads supporting the deletion are shown in an expanded view, with the same colors and supplementary alignments linked. Note, the same germline insertion appears narrower in the collapsed view than the expanded view, though both supporting and non-supporting reads are displayed along the same genomic coordinates. Supporting and non-supporting reads are categorized as such by the variant caller. **(F)** Protein structure translated from wild-type SHC2 transcript (REF) and from the transcript predicted to be formed by a mosaic exon 3 deletion (ALT). The protein formed from the ALT allele is missing the SH2 domain due to a premature stop codon and likely forms a substrate for nonsense mediated decay. See also Figure S14.

Consistent with their enrichment near genes, 8.3% (217 of 2610) of mosaic SVs directly disrupted exons and cis-regulatory coding elements, including distal enhancers (n=125), exons (n=61), proximal enhancers (n=31), CTCF-only elements (n=9), high-DNase-H3K4me3 elements (n=6) and promoters (n=7) (Figure 5B). Mosaic SV breakpoints were more common within exons than breakpoints of common germline SVs (p=0.0079) (Figure 5C), suggesting that gene-disrupting mosaic SVs may persist as their subclonal VAF limits deleterious impacts, whereas fitness compromising germline SVs undergo stronger negative selection.

To understand putative functional consequences of these mosaic variants, we applied a deep learning sequence-to-function model (AlphaGenome^64^) to all mosaic insertions (n=2071) and deletions (n=398), and a length and VNTR context matched set of Common Germline SVs from the same individuals. Mosaic deletions were predicted to interrupt 152 genome-wide splice sites as compared to 13 for the matched set of germline SVs (p=6.3×10^−5^), which could have significant downstream consequences from alternative splicing, non-functional transcripts, or nonsense mediated decay^65^ (Figure 5D). As an example, we observed a 514 bp mosaic deletion disrupting an exonic region of the SHC Adaptor Protein 2 (SHC2) gene (Figure 5E). This variant was predicted to lead to a novel splice junction between the 2^nd^ and 4^th^ exons of this gene, encoding a truncated protein lacking the SH2 domain that would likely be expressed at lower levels and be subject to nonsense mediated decay (Figures 5F and S14). The SHC gene family mediates multiple intracellular signaling pathways^66^, with possible associations between SHC2 alterations and adult-onset neurodegeneration^67^, highlighting potential deleterious impact of this variant that may in part be mitigated because it occurred at mosaic VAF. Together, the nonuniform distribution of mosaic SVs suggest that these alterations are likely to have substantial roles in generating functional variation across tissues and individuals.

### Mosaic variation across tissues in repeat element landscapes from short-read sequencing

To extend our observations of repeat element variation and tissue-specific mosaicism in settings where long-read sequencing may not be available, including from degraded or limited amounts of DNA (<10 ng), we applied ARTEMIS^45^, a kmer–based, alignment-free method that reconstructs genome-wide repeat element landscapes from conventional WGS (Figure 6A). By avoiding reference alignment, a major barrier to analyses of repetitive DNA in conventional short-read sequencing, ARTEMIS enables global quantification and cross-sample comparison of repeat element abundance. Although ARTEMIS does not identify specific SVs, we have shown that changes in kmer repeat landscapes correlated with the presence of diverse classes of SVs, including LINE-1 mediated deletions, gene amplifications, and complex breakpoints^45^. These observations suggested that ARTEMIS could be used to reveal tissue-specific patterns of mosaic variation.

**Figure 6.**
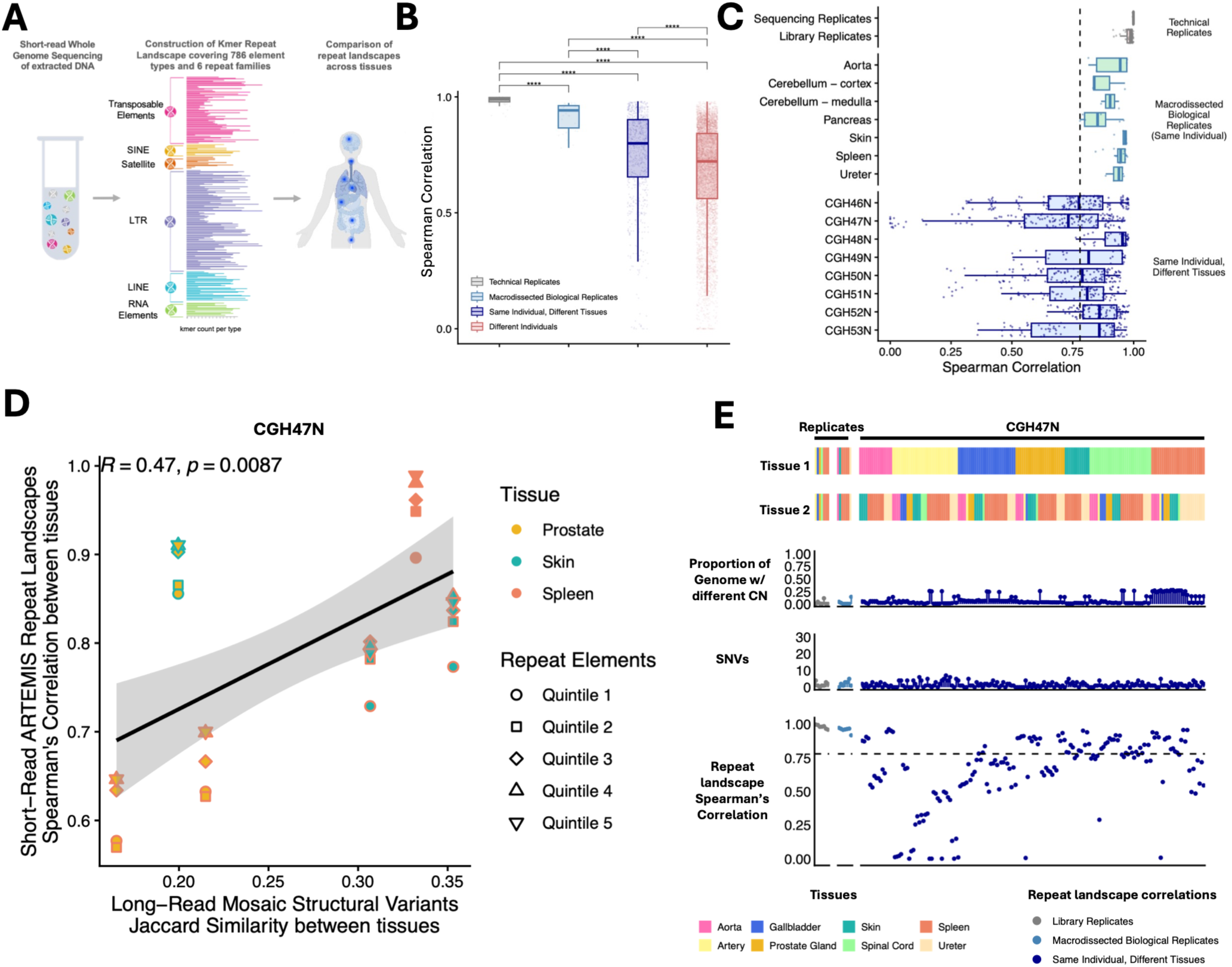
Intra-individual comparisons of global repeat landscapes constructed from short-read sequencing reveal widespread tissue-specific mosaicism across normal tissues. **(A)** Use of ARTEMIS alignment-free kmer repeat landscapes to reveal patterns of intra-individual mosaic variation. **(B)** Pairwise comparisons between kmer repeat landscapes show high concordance between repeat landscapes constructed from technical (sequencing or library) and biological (distinct tissue cores from the same organ) replicates, with decreased similarity between different organ tissues of the same individual, or between different unrelated individuals. Each point represents the median Spearman’s correlation coefficient across repeat element quintiles for a given comparison pair. **(C)** Increased variation in repeat landscapes across different tissues from the same individual is observed across all tissues and individuals studied. Each point represents the median Spearman’s correlation coefficient across repeat element quintiles for a given comparison pair. For individual CGH47N, the individual with the most distinct tissue types available for sequencing (n=8): **(D)** Tissue similarity as computed from short-read ARTEMIS repeat landscapes is concordant with similarity computed from long-read mosaic SVs. For each pair of tissues, the y-axis shows repeat landscape correlation, and the x-axis shows Jaccard similarity between mosaic SV calls. **(E)** For each pairwise tissue sample comparison, copy number differences, SNVs and repeat landscape correlations reveal tissue-specific repeat element differences to be the main driver of variation. See also Figures S15, S16 and S17.

We constructed ARTEMIS kmer repeat landscapes for 80 distinct tissue cores of 46 tissues from 8 individuals (range 4-8 tissue types per individual) sequenced with standard NGS (120 sequenced samples, including library and sequencing replicates) and performed a sample-specific loess correction to normalize for differences in sequencing coverage and duplicate rate (Figures 6A, S1A and S15A; Tables S11 and S12). We then computed pairwise spearman rank correlation coefficients between samples for each of the repeat element quintiles determined by expected genomic abundance, with lower correlation coefficients indicating less similarity in repeat landscapes between the compared samples (Figure S15B). Across repeat element groups, we observed high concordance between technical library or sequencing replicates (median pairwise correlation 0.99) and between biological replicates of the same tissue block (median pairwise correlation 0.94), while correlations dropped substantially for comparisons of different tissues within an individual (median pairwise correlation 0.80) and for comparisons across individuals (median pairwise correlation 0.72) (p<5.6×10^−14^ for all groups vs. technical replicates) (Figures 6B and 6C). These trends were consistent regardless of DNA library batch, operator, and sequencing flow cell (Figure S15C).

We next investigated whether patterns of repeat landscape mosaicism observed with ARTEMIS were similar to SV mosaicism identified via long-read sequencing. Analysis of individual CGH47N, for whom long and short-read sequencing data were available from the largest number of different tissue types (n=8 tissue types, n=13 samples, n=5 long-read libraries, n=10 short-read libraries) (Figures 1A and S1A) revealed high similarities between long-read mosaic SVs, and ARTEMIS repeat landscapes for each pair of samples (R=0.47, p=0.0087, Spearman’s Correlation) (Figure 6D). Comparison of long-read mosaic SVs and ARTEMIS between all pairs of samples from all tissue types showed a strong positive correlation between the approaches in 4 of 5 individuals analyzed (R=0.47, 0.69, 0.52 and 0.21 for CGH47N, 51N, 52N and 53N), with no obvious relationship in CGH50N, the individual with the least number of mosaic SVs identified (Figures 6D and S16A). Across individuals and tissues, pairwise ARTEMIS repeat landscape correlations varied, with related tissues showing high similarity (ie, Heart and Aorta, median correlation 0.97), and larger differences observed in tissues like spleen (median correlation 0.7) and pancreas (median correlation 0.82) that were also enriched for single-tissue mosaicism in the long-read SV analysis (Figures 6E, S16B and 3D).

To investigate whether ARTEMIS-detected repeat element variation could potentially be explained by copy number alterations (CNAs), we computed the percentage of the genome with different copy number changes between samples and found this to be generally less than 5% (median 1.2%, 1.3% and 2.0% for sequencing, library, and biological replicates, respectively, and 2.3% for comparisons of different tissues) consistent with prior reports of limited copy number variation within healthy tissues^68,69^ (Figures 6E, S16B). Overall, repeat landscape differences were not explained by CNAs or other genomic changes such as SNVs or small non-repeat element SVs, suggesting that postzygotic SV mosaicism within repeat elements is a widespread and previously underappreciated source of intra-individual genomic variation (Figures 6E, S16B, S17A and S17B).

### Repeat landscapes reflect both tissue-specific mosaicism and tumor-related changes

Lineage tracing for tissue-specific sources of genomic variation may also enhance our understanding of cancer. Tumorigenesis represents an extreme form of somatic variation, where alterations in a single cell of a tissue lineage give rise to a clonal population of tumor cells^13–15^. We hypothesized that if tumor-specific changes occur against a background of tissue-specific mosaicism, ARTEMIS repeat landscapes would capture both sets of changes (Figure 7A). We compared ARTEMIS repeat landscapes in 525 patients from the PCAWG (Pan Cancer Analysis of Whole Genomes) study^70^ comprising 12 types of primary solid tumors and their matched normal blood samples or normal adjacent tissues from where the tumor arose (n=1050 samples, Table S13). Repeat landscapes were more similar between tumors and normal adjacent tissue of the same type (n=121, median correlation 0.89) than when tumors were compared to a blood-derived normal (n=404, median correlation 0.86) (p=7.1×10^−7^ overall, p<0.05 for all tumor types with more than 10 samples with each type of normal, Wilcoxon’s rank-sum test) (Figures 7B and 7C). These tumor-related changes were comparable in scale to the extent of repeat element changes seen between different normal tissues of the same individual (median correlations across normal tissues ranged from 0.73 – 0.94 for the 8 individuals in Figure 6B).

**Figure 7.**
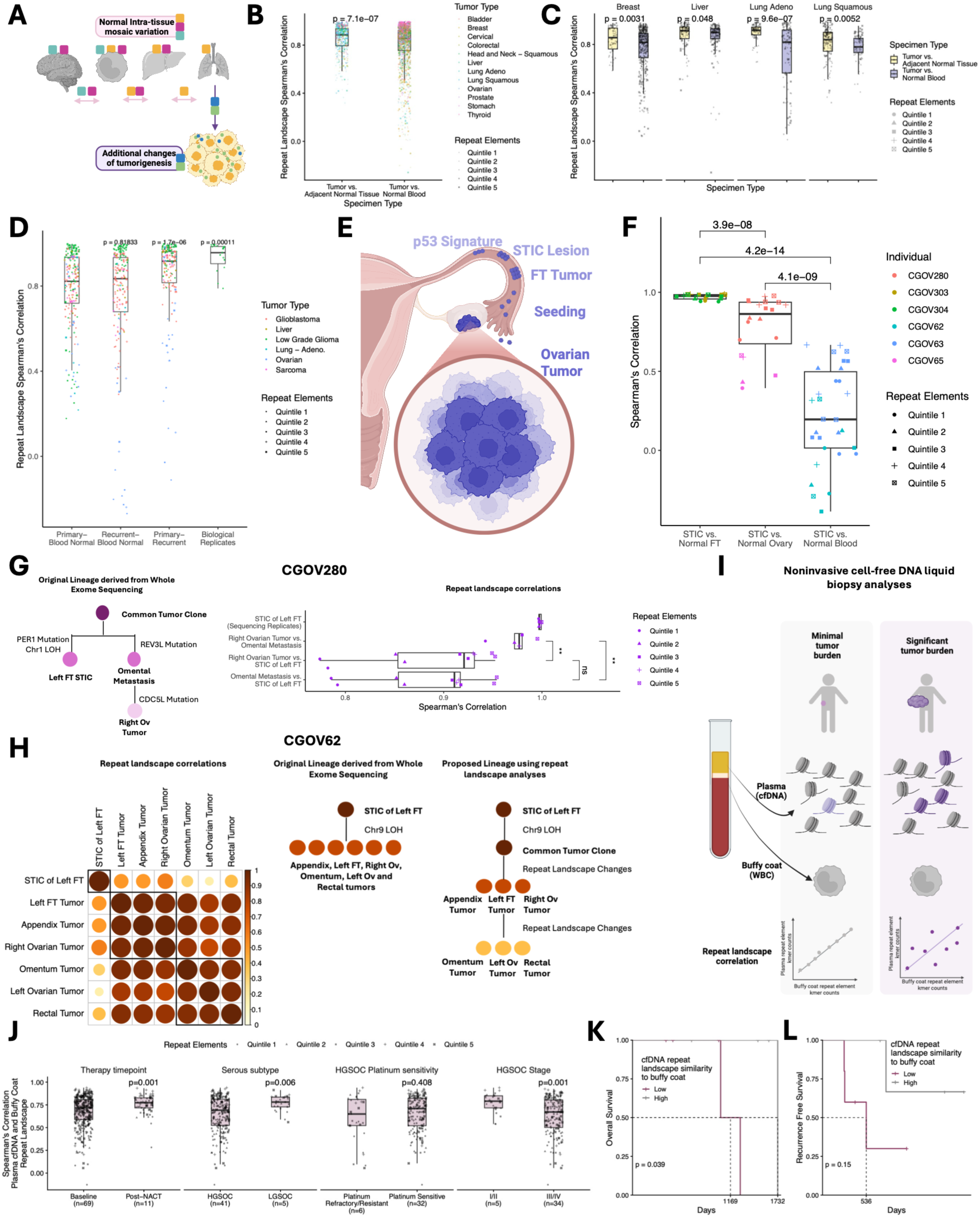
Tumor-specific and Tissue-specific changes in repeat landscapes characterize early tumorigenesis, illuminate fallopian tube origins of serous ovarian cancers, and enable noninvasive tumor monitoring in the blood. **(A)** Schematic of tissue-specific and tumor-specific changes. Mosaic variation is present between normal tissues of different types. During tumorigenesis, additional changes accumulate in cells originating from a single tissue type. **(B)** Comparisons between primary solid tumors and matched adjacent normal tissue or blood-derived normal demonstrate that tumor repeat landscapes are more similar to the normal tissue from which they originate than to blood. Each point represents the Spearman’s correlation coefficient for the given repeat element quintile and comparison pair. **(C)** Stratification of (B) by tumor type. All tumor types with at least 10 matched adjacent normal tissues are shown. **(D)** Comparisons between primary solid tumors, recurrent solid tumors from the same individual and matched blood-derived normals demonstrate that mosaic differences between blood and tumor are larger than differences between primary and recurrent tumors. Biological replicates are distinct samples of the same tumor. Each point represents the Spearman’s correlation coefficient for the given repeat element quintile and comparison pair. **(E)** Schematic illustrating fallopian tube (FT) origins of ovarian precancer lesions (ie, STICs, Serous Tubal Intraepithelial Carcinomas) that go on to seed ovarian tumors and peritoneal metastases. **(F)** Pairwise comparisons between kmer repeat landscapes show high concordance between repeat landscapes of STIC lesions and normal fallopian tube from the same individual, and progressively decreasing concordance between STIC and normal ovary, and STIC and blood, consistent with FT origins of STIC lesions. **(G)** Repeat landscape comparisons are consistent with evolutionary lineage originally derived from whole exome sequencing. Here, the left FT STIC is a distinct lesion from an omental metastasis that seeds a right ovarian tumor. Accordingly, repeat landscapes are highly concordant between sequencing replicates and between the right ovarian tumor and omental metastasis, but less concordant when tumors are compared to the distinct STIC lesion. **(H)** Repeat landscape correlations (left panel) illustrate a distinct left FT STIC, with highest similarity to a proximal cluster consisting of the left FT tumor, appendix tumor and right ovarian tumor, and more differences as compared to a distant peritoneal cluster of omental and rectal metastasis and a left ovarian tumor. The original whole exome sequencing lineage (middle panel) did not disambiguate between these 6 tumors arising from the left FT STIC, but repeat landscape analyses suggest accumulation of additional repeat landscape changes (right panel). **(I)** Schematic of how tumor- and tissue-specific changes to repeat landscapes may be noninvasively tracked using cell-free DNA liquid biopsies. Cell-free DNA comprises a mixture of DNA fragments from white blood cells, tumor cells, and normal organs. When tumor burden is low, repeat landscapes from cell-free DNA are expected to show high similarity to those from buffy coat (predominantly white blood cell), whereas when tumor burden is high more differences in the repeat landscape are expected. **(J)** Pairwise comparisons between kmer repeat landscapes from cell-free DNA and buffy coat of individuals with ovarian cancer associate with tumor characteristics and response to treatment. **(K)** After neoadjuvant chemotherapy, higher correlations between cfDNA and buffy coat are associated with longer overall survival, illustrating the potential for noninvasive tumor monitoring and characterization using measurements of repeat element mosaicism. **(L)** Similar to (K), showing that higher correlations between cfDNA and buffy coat after neoadjuvant chemotherapy trend with longer recurrence free survival. See also Figure S18.

We then extended this analysis to comparison of matched primary and recurrent tumors as well as blood derived normal cells from the same individuals (n=30 individuals, Table S14) from TCGA (The Cancer Genome Atlas). Here, both primary and recurrent tumors showed mosaic repeat differences from blood-derived normals (median correlations 0.82 and 0.84), but were significantly closer to each other (median correlation between primary and recurrent tumors 0.92) than to blood-derived normals (p<4.7×10-6, Wilcoxon’s rank-sum test) (Figure 7D). Overall, these observations suggest that in an individual patient, the normal tissue and blood cells are already separated by tissue-specific repeat element mosaicism, and these are further augmented by cancer-related alterations in the primary tumor, with minimal further changes in recurrent tumors.

### Repeat landscape mosaicism in ovarian pre-cancer lesions originating in the fallopian tubes

Since repeat landscapes capture mosaic changes in both normal tissue differentiation and subsequent tumor formation, we hypothesized that they may also reflect tissue of origin in pre-cancer lesions that could reveal a new class of pre-cancer alterations. High-grade serous ovarian cancer is thought to arise in the fallopian tube and subsequently seed the ovaries and the peritoneum^3^ (Figure 7E). Chromosomal copy number and mutational changes in ovarian tumors can be traced back to serous tubal intraepithelial carcinoma (STIC) lesions in the fallopian tubes^3^, but the minute DNA yield from these few-cell lesions has generally precluded long-read sequencing and limited the study of repeats and SVs in pre-cancer lesions. Using ARTEMIS, we traced lineage relationships among 30 samples of 24 laser-capture micro-dissected (LCMD) specimens^3^, including STICs (n=8 specimens), fallopian tube (FT) tumors (n=1 specimen), ovarian tumors (n=5 specimens), peritoneal metastases (n=4 specimens), and matched normal tissues (n=6 specimens) (Table S15).

Pairwise repeat landscape correlations were highest between STICs and matched normal fallopian tube epithelium (median 0.98), and lower for STICs versus normal ovary (median 0.86) and versus blood (median 0.20) (p < 3.9×10^−8^ for all comparisons, Wilcoxon’s rank-sum test) (Figure 7F). Prior lineage analysis using whole exome sequencing (WES) derived mutation and copy number analyses informed evolutionary relationships between FT STICs and subsequent FT, ovarian, and other patient tumors^3^ that were consistently reconstructed by ARTEMIS (Figure 7G). Analysis of changes in repeat elements through tumorigenesis using ARTEMIS further identified additional evolutionary substructures not evident through WES analyses alone, including among tumors that were previously thought to be identical in their lineage (Figure 7H). Together, these findings indicate that repeat landscapes capture tissue-specific signatures that can be used to refine lineage relationships among STICs, primary tumors, and metastases, complementing exome-based mutational and copy number phylogenies and illuminating early steps in serous tumor evolution.

### Tracking repeat landscape changes in cfDNA liquid biopsies for noninvasive tumor monitoring

Given our observations of tumor- and tissue-related repeat element changes among blood, cancer, and pre-cancer tissues, we examined whether these changes would be evident in cfDNA liquid biopsies that are beginning to be used for noninvasive detection and monitoring of cancer^71–80^ and other diseases^81^. cfDNA of nucleosomal origin is released in the circulation from dying cells, and arises primarily from white blood cells, though aberrant tumor tissue-derived contributions are also found in cancer^72,74,82,83^. We have previously shown that genome-wide repeat landscapes constructed from low coverage WGS of the compendium of cfDNA fragments can be used for early detection of lung, liver and other cancers^45^. Here, we compared repeat landscapes from plasma cfDNA and buffy coat genomic DNA for 72 individuals with ovarian cancer, collected prior to treatment (n=69) and after resection and neoadjuvant chemotherapy (NACT) (n=11) (Table S16).

We hypothesized that cfDNA liquid biopsies with higher tumor burdens would be less similar to buffy coat, as their repeat landscape would reflect both differences from normal tissue-mosaicism as well as tumor-related changes in cancer cells (Figure 7I). Accordingly, we observed that correlations between ARTEMIS repeat landscapes from cfDNA and buffy coat were higher post-NACT as compared to the pre-treatment baseline samples, consistent with a decrease in ovarian tumor-derived cfDNA following treatment (p=0.001, Wilcoxon’s rank-sum test) (Figure 7J). Similarly, correlations between cfDNA and buffy coat were higher in low-grade and early-stage tumors, and trended higher in platinum-sensitive HGSOC as compared to platinum-resistant HGSOC, consistent with lower circulating tumor fractions or reduced burden of somatic genomic changes in earlier or less aggressive tumors (p=0.006 for LGSOC vs. HGSOC and p=0.001 for stages I/II vs. III/IV HGSOC, Wilcoxon’s signed rank-sum test) (Figure 7J). After NACT, individuals with higher correlation between cfDNA and buffy coat repeat landscapes had significantly longer overall survival (median 4.75 years vs. 3.2 years for individuals with correlations above vs. below the median, p<0.039) and similar trends for recurrence-free survival (median not reached vs. 1.5 years for individuals with correlations above vs. below the median) (Figures 7K, 7L and S18), highlighting the ability for noninvasive analyses of repeat landscape mosaicism to capture response to therapy.

## DISCUSSION

This study provides evidence that mosaic SVs are prevalent across normal human development and healthy life, and that similar tumor-related changes occur in tumorigenesis. By leveraging multi-tissue long-read sequencing, we identified mosaic SVs as significant drivers of variation among tissues in an individual. Mosaic SVs impacted a median 285.2 Kb per individual across the genome and 8.3% of these alterations were predicted to disrupt functional genomic elements, with consequences for transcript splicing and protein structure. These SVs were primarily in genomic repeats, which are among the most variable regions in the germline and are known to exhibit error-prone replication^31,32,48,49^. We found that repeat-mediated mechanisms, especially L1-driven retrotransposition and replication-based template switching, are the predominant driver of mosaic SVs. This structural mosaicism is non-uniformly distributed across the genome, and points to a previously unappreciated^3,7,8,16–19^ role for these alterations in generating functional variation across both tissues and individuals.

To extend observations of structural mosaicism using accessible short-read sequencing approaches, we adapted ARTEMIS^45^, an alignment-free, kmer–based approach that quantifies genome-wide repeat landscapes, to identify tissue-specific mosaic changes in repeat elements. Beyond providing complementary support for our long-read findings, these repeat landscape signatures enabled study of similar changes in tumorigenesis. We identified larger differences between tumors and blood-derived normal samples than adjacent normal tissue samples, pointing to both tissue-specific and tumor-specific repeat element alterations. Analysis of repeat landscape changes enabled lineage tracing in pre-cancer samples, an analysis previously limited to SNVs, small indels, and CNAs, and enabled non-invasive blood-based detection of tumor SVs as well as identification of such alterations during therapy. These findings highlighted the tissue-specific origins of early ovarian tumorigenesis and highlighted repeat element changes as an additional class of somatic alteration that may be used to refine our understanding of tumor evolution.

This study has some limitations. First, detection of low VAF SVs remains technically challenging^21–23,25^. To minimize the chances of artifactual calls, we incorporated extensive replicates into the study design including both technical (sequencing and library) and biological (distinct DNA samples), analyzed mosaic benchmark^21^ and trio sequencing^49^ data for ground truth error rates in mosaic calls, implemented variant confidence filters, performed modeling of technical missingness in SV calls, and developed a short-read kmer based approach to orthogonally validate these large, repeat rich SVs with a separate sequencing chemistry. Regardless, some tissue-restricted patterns may persist due to limited sensitivity, and breakpoint identification remains challenging, particularly within VNTRs. A consequence of our filtering and moderate sequence coverage is that some ultra-low VAF calls were likely missed, and therefore we have likely underestimated the number of true mosaic calls. Nevertheless, our approach provides a balance between biological discovery of previously underappreciated mosaicism while limiting artifactual calls. A second limitation is that the short-read sequence analyses we performed were correlative and, unlike long-read sequencing, could not identify specific mechanisms of detected repeat element alterations. However, the consistency of mosaic signatures from long-read and short-read sequencing data suggests that these events are widespread and opens the door to their global assessment in short-read sequencing of limited input samples, and in fragmented cfDNA for noninvasive tumor monitoring in the blood.

Future studies in larger cohorts with deeper sequencing coverage will be needed to further elucidate mechanisms and functional impacts of mosaic SVs and to refine estimates of their prevalence and developmental timing. Nonetheless, this study presents multi-tissue evidence of widespread postzygotic mosaicism in SVs, characterizes their functional consequences, and establishes signatures of previously underappreciated changes in repeat elements as a useful measure for lineage tracing in normal development and tumorigenesis.

## MATERIALS AND METHODS

### Study populations

The JHU cohort consisted of eight individuals without cancer, with fresh-frozen tissue blocks (n=47) purchased from BIOIVT (Tables S1 and S2). The tissue blocks encompassed 21 different organ types with a range of 4-8 types per individual.

We additionally analyzed long-read sequencing data from 7 individuals sequenced by the Genome in a Bottle Consortium (GIAB)^49^, including two parent off-spring trios. For each individual two library replicates were constructed from blood cell DNA and sequenced via Oxford Nanopore. We further obtained *in silico* benchmarking ONT sequencing data constructed from mixtures of DNA from distinct individuals. This SMaHT-MIMS benchmark is designed to simulate low VAF mosaic structural variants at 30X, 60X and 90X coverage^21^.

For analyses of tumors and normal tissues, we selected all 1078 PCAWG samples (539 matched Tumor/Normal pairs) from 12 tumor types in the Pan-Cancer Analysis of Whole Genomes (PCAWG)^70^, and excluded 14 pairs on the PCAWG blacklist, leaving 1050 samples consisting of 525 tumor/normal pairs (breast (n=91), lung (n=86), colorectal (n=60), liver (n=54), thyroid (n=48), head and neck squamous cell (n=44), ovarian (n=42), gastric (n=38), bladder (n=23), cervical (n=20) and prostate (n=19)). For these 525 tumors, matched normal tissues were either blood-derived (n=404) or normal adjacent tissue (n=121) (Table S13). We additionally selected 30 recurrent tumors from The Cancer Genome Atlas (TCGA), and identified additional samples from those individuals including blood-derived normal, primary tumor, and biological replicates for tumor tissue (Table S14). For this recurrent tumor analysis, we restricted to samples sequenced at the same site and with the same read length in order to minimize technical variability.

For analyses of pre-cancer lesions, ovarian tumors, peritoneal metastases and associated normal tissues we performed whole genome sequencing (n=30) on DNA libraries (n=26, plus 4 sequencing replicates) constructed from 24 laser-capture micro-dissected (LCMD) specimens, including STICs (n = 8 specimens), fallopian tube tumors (n = 1 specimen), ovarian tumors (n = 5 specimens), peritoneal metastases (n = 4 specimens), and matched normal tissues (n = 6 specimens) (Table S15). These libraries were originally used for targeted capture and exome sequencing in a prior study^3^; here we utilized remaining pre-capture library material.

For whole genome sequencing analyses of cfDNA and genomic DNA from matched buffy coat for individuals with ovarian cancer (n=72), we utilized plasma and buffy coat separated from whole blood samples collected via venipuncture at the University of Minnesota, Division of Gynecologic Oncology (Table S16).

### Macrodissection of normal tissues

For the JHU Cohort, we embedded each tissue sample in OCT within an embedding mold. For most tissues, we cut an H&E slide and annotated histologically similar regions for two sets of tissue cores to be used as biological replicate samples. We then positioned the slide alongside the OCT block and collected two distinct sets of tissue needle cores which were then processed separately as biological replicates. Where histology was not available, we collected a single set of tissue needle cores, and after DNA extraction split for preparation of 2 DNA libraries (library replicates). We extracted genomic DNA using the Qiagen QIAamp DNA FFPE Tissue kit. We quantified extracted DNA yield via Qubit.

### Long-read sequencing and structural variant calling

We constructed libraries for long-read sequencing using up to 3 ug of input DNA with the Ligation sequencing DNA V14 protocol (SQK-LSK114, Oxford Nanopore Technologies), with the following modifications to the manufacturer’s instructions: (i) Incubate for 8 minutes rather than 5 minutes during all rotator mixer incubations (ii) Incubate for 5 minutes rather than 2 minutes during DNA repair and end-prep DNA recovery from beads (iii) Incubate for 18 minutes rather than 10 minutes during adapter ligation and clean-up DNA recovery from beads.

Library concentrations were quantified by Qubit, and where possible 300ng of library was loaded on a FLO-PRO114M flow cell. Sequencing was performed on a PromethION sequencer with the following software versions: MinKNOW: 23.11.7, Bream: 7.8.2, Configuration: 5.8.6, Dorado: 7.2.13, MinKNOW Core: 5.8.6.

For individuals CGH47N, CGH50N and CGH51N, for libraries with more than 300ng available, we performed a re-load with available material after washing the flow cell every 24 hours, over a total of 72 hours. For individuals CGH52N and CGH53N we loaded all library up to a maximum of 300ng and ran for 72 hours without reload. If more than 300ng of library was available, we ran a second flow-cell and then combined sequencing output for downstream analyses.

We basecalled all POD5 using the Dorado sup@v.5.0.0 model (https://github.com/nanoporetech/dorado). We aligned sequencing reads with Minimap2 -ax lr:hq preset^84^, sorted and indexed bams with Samtools^85^, and performed phasing and haplotyping with Clair3^86^ and LongPhase^87^. We then performed SV calling with Severus^37^. For second caller SV confirmation, we used Sniffles2^24^.

### Categorization of SVs

We first filtered to a high-quality set of variant calls (MAPQ > 50) and required that in at least one of the tissue samples for the individual in which the variant was called, 20+ reads spanned the breakpoint. We also excluded any SVs occurring on ChrY in female individuals.

We then classified variants as follows:

1. Common Germline: If a variant occurred in the 1000 Genomes long-read call set^48^, or within 10bp of a different variant in 1000 Genomes, we classified it as Common Germline.
2. We excluded any variants called in multiple unrelated individuals that were not found within 1000 Genomes, as we hypothesized these were likely artifactual calls in hard to resolve regions of the genome.
3. Rare Germline or *De Novo*: Among the remaining variants, if the SV occurred at VAF > 0.25 in any tissue of the individual in which the variant was called, we classified it as Rare Germline or *De Novo*.
4. Mosaic: The remaining variants occurred at VAF ≤ 0.25 in all tissues in which they were found and were classified as mosaic
5. For variant clusters and complex SVs with ambiguous classification, we categorized all SVs belonging to the cluster as follows:

a. Common Germline: If any portion of the cluster occurred in 1000 Genomes, we classified all SVs in the cluster as Common Germline
b. Rare Germline or *De Novo*: If any portion of the cluster occurred at VAF > 0.25 in any tissue, but not in 1000 Genomes, we classified all SVs in the cluster as Rare Germline or *De Novo*.
c. Rare Germline or *De Novo*: If multiple SVs were called at a given location for an individual, and any of those SVs occurred at germline VAF (>0.25) we designated all variants at that location as Rare Germline or *De Novo*.

We annotated insertion and deletion SV calls for repeat-element involvement with SVAN (https://github.com/REPBIO-LAB/SVAN) (Tables S9 and S10). We also called indels (<50 bp) and single nucleotide variants (SNVs) in the GIAB data using clair3^86^ and applied the above classification scheme using the previously published 1000 Genomes indel/SNV call set^88^.

### Benchmarking of SV classification scheme using Genome in a Bottle and SMaHT-MIMS reference data

We applied the SV calling and classification approach described above to the 30X, 60X and 90X ONT sequencing data from the SMaHT-MIMS benchmark DNA mixtures. We quantified recall (sensitivity) as the proportion of true SVs in the benchmark that were identified by our approach, both before and after confidence filtering. We quantified precision as the proportion of SV calls we made that corresponded to SVs within the benchmark set.

To estimate the true burden of mosaic SVs in the JHU Cohort, we performed a Monte Carlo simulation incorporating benchmark-derived uncertainty in precision and recall. For each VAF bin, precision and recall were modeled as Beta distributions parameterized by the number of true positives, false positives, and false negatives observed in the benchmark dataset. For each of 10000 iterations, precision and recall values were sampled from their respective Beta distributions, and the total number of true SVs was estimated as the number of observed SV calls multiplied by the sampled precision and divided by the sampled recall. This procedure generated a distribution of plausible true SV counts per VAF bin, from which we report the median and standard deviation.

We additionally analyzed ONT sequencing data from two parent-offspring trios within GIAB and implemented our SV calling and classification approach above. We used parental sequencing data to distinguish between rare germline and *de novo* SVs and quantify the rate of *de novo* SVs. We also quantified rates of two types of artifactual calls that could occur using our approach. First, among putative mosaic SVs, we defined false positive mosaic calls as those that were found in the offspring at subclonal VAF but also confirmed in the parent. Second, among common germline variants, we defined false negative calls as those found in the offspring and in 1000 Genomes but not found in either parent.

### Long-read cell-type methylation deconvolution in normal tissues

We constructed a reference atlas from whole genome bisulfite sequencing (WGBS) data of sorted normal primary cells^82^ aligned to hg19. For each sample in the atlas, beta values were computed at each CpG as the ratio of methylated reads to total coverage. We selected cell types corresponding to blood, fibroblast, smooth muscle, endothelium and any cell types derived from the tissues we studied (Aorta, Spleen, Cerebellum, Pancreas, Prostate, Skin). Cell types were condensed by shared lineage into meta-groupings as follows: T cells [total CD3, CD4, and CD8], monocytes and macrophages [blood monocytes; colon, liver, lung alveolar, and lung interstitial macrophages], smooth muscle [aorta, coronary artery, bladder, and prostate], vascular endothelium [aorta and saphenous vein], endothelium [lung alveolar, kidney glomerular, kidney tubular, pancreas, and pancreatic islet], and neurons [cortex and cerebellum]. Granulocytes, B cells, NK cells, dermal fibroblasts, oligodendrocytes, epidermal keratinocytes, prostate epithelium and the five pancreatic cell types [acinar, duct, alpha, beta, delta] were retained as individual references, for a total of 18 reference groups. Per-CpG methylation for each reference group was computed as the mean beta value across all constituent samples. These groupings were guided by the hierarchical clustering reported for the source atlas^82^. At each CpG site, we computed an F statistic comparing between-group to within-group variance in beta values across the 18 reference groups. The top 5% of CpG sites with largest F statistic (∼1.4 million) were retained as markers.

In our JHU Cohort ONT samples, we extracted per-CpG methylation using Modkit with the pileup function, aggregating 5mC and 5hmC modifications across both strands to match the bisulfite-derived reference, which does not resolve the two modifications. Cell-of-origin proportions were estimated at the marker CpGs with MethID^89^, which fits sample methylation as a non-negative, sum-to-one mixture of the reference profiles. Marker CpGs covered by fewer than two reads in a sample were excluded from the fit. We did not perform deconvolution on our JHU Cohort ureter samples as no ureter cell types were available in the published atlas. For visualization, we aggregated the 18 reference groups as follows: Blood [T cells, monocytes and macrophages, granulocytes, B cells, NK cells], Brain cells [neurons, oligodendrocytes], endothelium, vascular endothelium, fibroblasts [dermal fibroblasts], pancreas cells [acinar, duct, alpha, beta, delta], prostate cells [prostate epithelium], skin cells [epidermal keratinocytes], and smooth muscle.

### Orthogonal validation of long-read called mosaic SVs using short-read sequencing

Since large, repeat-involved structural variants cannot be resolved in conventional short-read sequencing, we developed an alignment-free kmer based approach to orthogonally validate mosaic SVs in the JHU Cohort. We identified all 24bp sequences (kmers) that spanned a mosaic SV breakpoint. The kmers not found elsewhere in the chm13 reference were considered novel mosaic breakpoint kmers. To be included in this set, a kmer had to span the breakpoint, meaning that atleast 1bp had to be found on either side of the breakpoint. We also considered subsets of these novel kmers with minimum flanks of 6bp or 12bp on either side of the breakpoint. We counted these novel kmers in all short-read sequencing reads from samples of the individual with the SV and unrelated individuals, and normalized the counts for sequencing coverage. We considered a mosaic SV to be evaluable if atleast 1 novel breakpoint kmer was found in atleast 1 sample. We considered an evaluable SV to be Validated-Enriched if the median kmer count for that SV was higher in the individual with the variant than unrelated individuals. We considered an evaluable SV to be Validated-Exclusive if the median kmer count in unrelated individuals was 0.

### Simulations of technical missingness in mosaic variant calling

For mosaic variants not called in all tissue samples, we sought to determine whether these represented technical missingness (ie, due to coverage) or true biology (ie, lineage restricted clones). The following procedure was performed for each individual in the JHU Cohort separately (CGH47N, CGH50N, CGH51N, CGH52N, CGH53N).

For each mosaic SV in each tissue sample, we recorded alternate-allele read count (*ALT*) and reference-allele read count (*REF*), with total depth *DP=ALT+REF*. We also recorded *t*, the tissue type (ie, spleen, prostate etc); *v*, the unique variant ID; and *i*, the tissue sample (ie, spleen – 1, spleen – 2).

We fit a beta-binomial generalized linear mixed model with a logit link for the *ALT* fraction with fixed effects for *base_tissue* and random intercepts for *var* and *sample_id*:

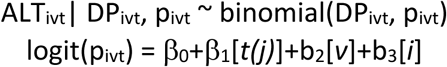

Here, β_0_ is an intercept, β_1_[*t(j)*] are fixed effects for tissue type, and b_2_[*v*] and b_3_[*i*] are random intercepts for variant and tissue sample. The fixed effects for the base tissue type estimate tissue-specific shifts in the underlying *ALT* fraction (ie, detectability differences due to tissue cellularity). The random intercept for variant captures heterogeneity between different variants in the callset (ie, due to clone size). The random intercept for tissue sample captures sample-level sequencing quality differences (ie, coverage).

The model was fit by maximum likelihood using the R package *glmmTMB* ^90^. From the fitted model we obtained, for each variant, *v*, and tissue sample, *i*, of base tissue type, *t*, the expected *ALT* fraction p^_ivt_. Therefore, the probability of detection is the probability of at least one read carrying the variant in DP_ivt_=ALT_ivt_+REF_ivt_ trials with binomial probability p^_ivt_. Formally:

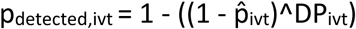

After computing these probabilities for every variant and tissue sample, we summarized tissue-level detection probability for each variant as the union across all samples in that tissue. For example, if there are r=3 samples of a given tissue, *t*, we would summarize detection probability for variant *v* as:

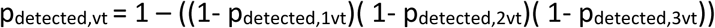

We then performed 1000 simulation trials as follows. For one trial:

- draw tissue presence for each variant, *v*, in each tissue, *t*, as Bernoulli(p_detected,vt_)
- across all variants, summarize the counts for each mosaic signature. For example, a signature may consist of “Present in Tissue 1 and Tissue 2, not present in Tissue 3”.

After the 1000 simulations, for each signature we defined the expected variant count as the mean number of occurrences across trials.

Then, for each signature we estimated an empirical one-sided p-value as the fraction of simulations with simulated counts ≥ the observed (using a +1 correction in the numerator and denominator to avoid p-values below the theoretical precision of our simulation). To account for multiple signatures, we used Benjamini–Hochberg corrected p-values. We defined enriched signatures (ie, those representing biology as opposed to technical missingness) as those with a corrected p-value < 0.05.

### Characterizing functional significance of mosaic SVs

We examined the relationship between SVs that we called in the JHU cohort and various genomic features: replication timing, GC content, distance to gene body, distance to exon, distance to centromere, and distance to telomere. We binned the chm13 genome into 1000bp bins. For each bin, we computed the proportion of guanine and cytosine bases and divided bins into 5 quintiles from low to high GC content. We obtained centromeric coordinates for the chm13 assembly^47^ and computed the distance between each chromosome’s centromere and the nearest edge of each bin on that chromosome, and divided bins into quintiles from near to far. Bins overlapping the centromere were considered to have a distance of zero. We repeated this procedure for telomeres. For gene bodies, we obtained coordinates for combined intronic and exonic regions for each gene from the NCBI RefSeq chm13 track and computed the distance between the edges of bins and the nearest gene. For bins overlapping a gene we set the distance to zero. We then divided bins into quintiles from near to far. We repeated this procedure using only exon intervals. For replication timing, we downloaded replication timing tracks^91,92^ for the IMR90, HNEK and GM12878 cell lines from the UCSC Genome Browser. These tracks were computed by averaging the wavelet-smoothed transform of the six fraction profile, representing different time points during replication in 1 kb bins. We computed the weighted average in each bin, with higher values indicating earlier replication timing, lifted coordinates over from hg19 to chm13 and divided the bins into quintiles from early to late replicating. We then assigned each SV to a quintile for each feature based on the genomic bin the SV overlapped. For large SVs overlapping multiple bins, or for inter- and intra-chromosomal SVs with multiple sets of breakpoints, we assigned the SV based on the bin with the largest overlap.

We also examined overlap of SVs with the actual genomic coordinates of functional elements including gene bodies (exons and introns) defined by the NCBI RefSeq track, and cis-regulatory elements defined by the ENCODE consortium^93^, with coordinates lifted over from hg38 to chm13. We grouped cis-regulatory elements defined by ENCODE as follows: Distal enhancers (dELS and dELS, CTCF-bound regions defined by ENCODE), proximal enhancers (pELS and pELS, CTCF-bound), promoters (PLS and PLS, CTCF-bound), high-DNase-H3K4me3 elements (DNase-H3K4me3 and DNase-H3K4me, CTCF-bound), and CTCF-only elements (CTCF-only, CTCF-bound).

### Structural variant effect prediction using AlphaGenome

For each mosaic SV, matched reference (REF) and alternate (ALT) 1,048,576-bp sequence windows were constructed from the chm13 assembly and scored directly with AlphaGenome’s predict_sequence() interface. REF windows were centered on the variant position where possible unless near a chromosome end in which case the window was slid toward the SV to accommodate the full length. ALT windows were built by removing (deletions) or inserting (insertions) the variant-length sequence at the breakpoint, with the upstream flank held identical between REF and ALT to avoid coordinate-realignment artifacts. Predictions were generated for expression and splicing tracks covering the UBERON ontology terms corresponding to aorta, pancreas, cerebellum, skin, prostate, ureter and spleen tissues. For each mosaic SV, we selected a common germline SV as a comparator drawn from the same individual, matched on SV length and VNTR-overlap status. We also generated variant scoring predictions for the matched set of germline SVs. Splice donor and acceptor probability scores (from the SPLICE_SITES track) were extracted from REF and ALT at every position overlapping each variant. A site was called disrupted if its REF score exceeded 0.1 and fell below 0.001 in the corresponding ALT prediction.

For the 514-bp mosaic deletion overlapping an exon of *SHC2*, the REF and ALT contigs were built and scored as above using the SPLICE_SITES and SPLICE_JUNCTIONS tracks. Loss of both splice sites flanking the affected exon, together with a predicted novel junction directly joining its flanking exons, was interpreted as exon skipping. The resulting frameshift relative to annotated *SHC2* transcripts was used to determine the new amino acid sequence of the protein generated by the ALT allele. The REF and ALT amino acid sequence was scored using AlphaFold to predict the truncated protein structure generated by the ALT allele and per-residue confidence scores were assessed across each known protein domain.

### Visualization of mosaic SVs in the JHU Cohort

For each tissue sample and mosaic SV, we extracted all reads overlapping the breakpoint and a +/− 1000bp flank, and merged the reads into a single bam file, with the tissue sample indicated in the read group (RG) field. We then separated reads identified as “supporting” by Severus, the long-read variant caller^37^. We then visualized both the supporting reads bam and the non-supporting reads bam using IGV (Integrated Genomics Viewer), with reads colored and sorted by read group (tissue sample, in our case).

### CRISPR cell-line analysis of impacts of double stranded DNA breaks within repeat elements

We downloaded data from the DepMap portal (https://depmap.org/portal) for a CRISPR screen of 318 cell lines using Sanger’s KY Cas9 library^57^. For each of 89,432 19bp sgRNAs, we used Meryl^94^ to count the number of instances of the sgRNA sequence and its reverse complement. We also determined how many of those instances overlapped a repeat element as defined by the chm13 T2T repeat masker track, encompassing transposable elements, human satellites, long and short interspersed repeat elements (LINEs and SINEs), long terminal repeats (LTRs) and other classes of repeat elements. For each sgRNA and cell line, we computed an adjusted log-fold change (*a*) as the log2-fold-change for pDNA counts in the screen as compared to the original plasmid pool minus a constant “median non-essential depletion” computed for each cell line and provided by the DepMap consortium. Previous studies have shown that sgRNA introduction of double stranded breaks (DSBs) have deleterious impacts on cell fitness independent of gene function ^95^.

For each cell line screen, we fit a linear model for *a*, the adjusted log fold change:

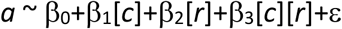

Here, *c* is the log1p transformed number of occurrences of each sgRNA within the chm13 reference genome, *r* is the log1p transformed number of occurrences of each sgRNA within annotated repeat elements, and ε is the independent residual errors.

Since a negative adjusted log fold change corresponds to a deleterious impact on cell survival, we interpreted negative values of β_1_ and β_2_ as evidence of deleterious impacts of additional DSBs introduced by the sgRNA sequence, and a positive coefficient, β_3_, for the interaction term to suggest that when these additional DSBs occurred within repeats the deleterious impact was attenuated.

### Short-read sequencing and generation of repeat landscapes

For the JHU Cohort, we sheared genomic DNA in microtubes using the truXTRAC™ FFPE DNA Kit (Covaris) with 5 minute shearing time for 100ng shearing input and 10 minute shearing time for 1ug shearing input and quantified DNA via Agilent Tapestation. We then prepared whole genome sequencing libraries with 15ng of input DNA and 4 cycles of PCR amplification using the NEBNext DNA Library Prep Kit for Illumina (New England Biolab) with four main modifications to the manufacturer’s guidelines: (i) The library purification steps followed the on-bead AMPure XP (Beckman Coulter) approach to minimize sample loss during elution and tube transfer steps; (ii) NEBNext End Repair, A-tailing, and adapter ligation enzyme and buffer volumes were adjusted as appropriate to accommodate on-bead AMPure XP purification; (iii) Illumina dual index adapters were used in the ligation reaction; and (iv) Libraries were amplified with Phusion Hot Start Polymerase. We sequenced 100bp paired-end reads on the Illumina NovaSeq 6000. Previously constructed pre-capture DNA libraries for pre-cancer lesions, ovarian tumors, peritoneal metastasis and associated normal tissues ^3^ were quantified via Agilent Tapestation and sequenced with 100bp paired-end reads on the Illumina NovaSeq 6000. We also obtained paired-end sequencing data for n=1050 PCAWG and n=124 TCGA samples consisting of primary tumors, recurrent tumors and tissue adjacent and blood derived normals.

We aligned sequenced samples using Bowtie2^96^, and identified copy number alterations using CNVKit^97^, and SNVs and small indels (<50bp) using Manta^98^ and Strelka^99^. We constructed alignment-free kmer repeat landscapes using the ARTEMIS pipeline (https://github.com/cancer-genomics/artemis_pipeline) we previously described ^45^. Briefly, in unaligned sequencing reads, we counted 1.2 billion kmers that uniquely identify one of 1280 distinct repeat element types within SINE, LINE, LTR, human satellite and transposable element families. We then selected kmer counts for the 786 repeat elements with an expected abundance of at least 1000 kmers/million aligned reads. To correct for differences in sequencing coverage, duplicates and other technical differences between samples, we performed a loess regression on expected kmer abundance based on the chm13 reference genome vs. observed counts for each sample, and defined the adjusted count as the ratio of observed count to count predicted by the sample-specific loess model. To compare these adjusted kmer repeat landscapes between individuals, we stratified the 786 repeat elements into quintiles, and computed a Spearman correlation coefficient for each quintile. Lower correlation coefficients were interpreted as representing larger differences in the repeat landscape.

To test whether long-read-derived mosaic SV sharing between tissues reflects the same underlying tissue relatedness captured by ARTEMIS repeat-landscape correlations, we computed the Jaccard similarity between mosaic SV calls for each pair of samples (the number of mosaic SVs detected in both samples divided by the number detected in either sample). We then compared Jaccard similarity between long-read mosaic SV profiles for a sample pair to the corresponding ARTEMIS short-read repeat landscape correlation. Concordance between the two metrics was assessed using Spearman rank correlation across all sample pairs within an individual.

### Noninvasive tumor monitoring using genome-wide repeat landscapes constructed from blood and cfDNA samples

We extracted cfDNA plasma cfDNA and genomic DNA from buffy coat for 72 individuals with ovarian cancer, collected at baseline prior to treatment (n=69) and after resection and neoadjuvant chemotherapy (NACT) (n=11), and prepared whole genome sequencing libraries for illumina sequencing as described above and in prior publications ^71,100^. We computed kmer repeat landscapes and pairwise correlations between samples as described above using the ARTEMIS methodology^45^. We evaluated these correlations between cfDNA and buffy coat for each individual with respect to clinical variables including treatment timepoint, tumor subtype and stage, recurrence free survival, overall survival and sensitivity to platinum-based chemotherapy.

### AI Usage Statement

For some analyses, gpt-5, gpt-5.2, and claude-sonnet-4.5 were used to generate initial versions of code snippets for bash and R based on descriptions of required inputs, functionality and outputs from the authors. All LLM-generated code was verified and tested by the authors and revised as needed for manual integration with other author-derived scripts and pipelines. No end-to-end AI generated code was used to generate figures, results, or conclusions.

## Acknowledgements

We thank members of our laboratories for critical review of the manuscript. Results shown here are in part based upon data generated by the Genome in a Bottle Consortium, the 1000 Genomes Project, the Telomere to Telomere (T2T) Consortium, the Somatic Mosaicism Across Human Tissues (SMaHT) consortium, the TCGA Research Network, and the DepMap Consortium.

## Data and Code Availability

Code to generate ARTEMIS repeat landscapes from WGS FASTQ files may be found at https://github.com/cancer-genomics/artemis_pipeline. Code to run the Severus long-read SV calling pipeline may be found at https://github.com/KolmogorovLab/Severus. Code to run the SVAN repeat element annotation pipeline for SV calls may be found at https://github.com/REPBIO-LAB/SVAN/tree/main. Code and instructions for querying the AlphaGenome API may be found at https://github.com/google-deepmind/alphagenome. The AlphaFold server to query amino acid sequences may be found at https://alphafoldserver.com/welcome. Sequence data generated for samples in this study will be deposited in the European Genome-Phenome Archive (EGA) under EGAS00001008557 and EGAS50000001331 as permitted by institutional regulations. PCAWG/TCGA BAM files were downloaded from Bionimbus (icgc.bionimbus.org). The Bionimbus and EGA data are available through controlled access and have restricted use to approved investigators.

## Supplementary Figures

**Figure S1.**
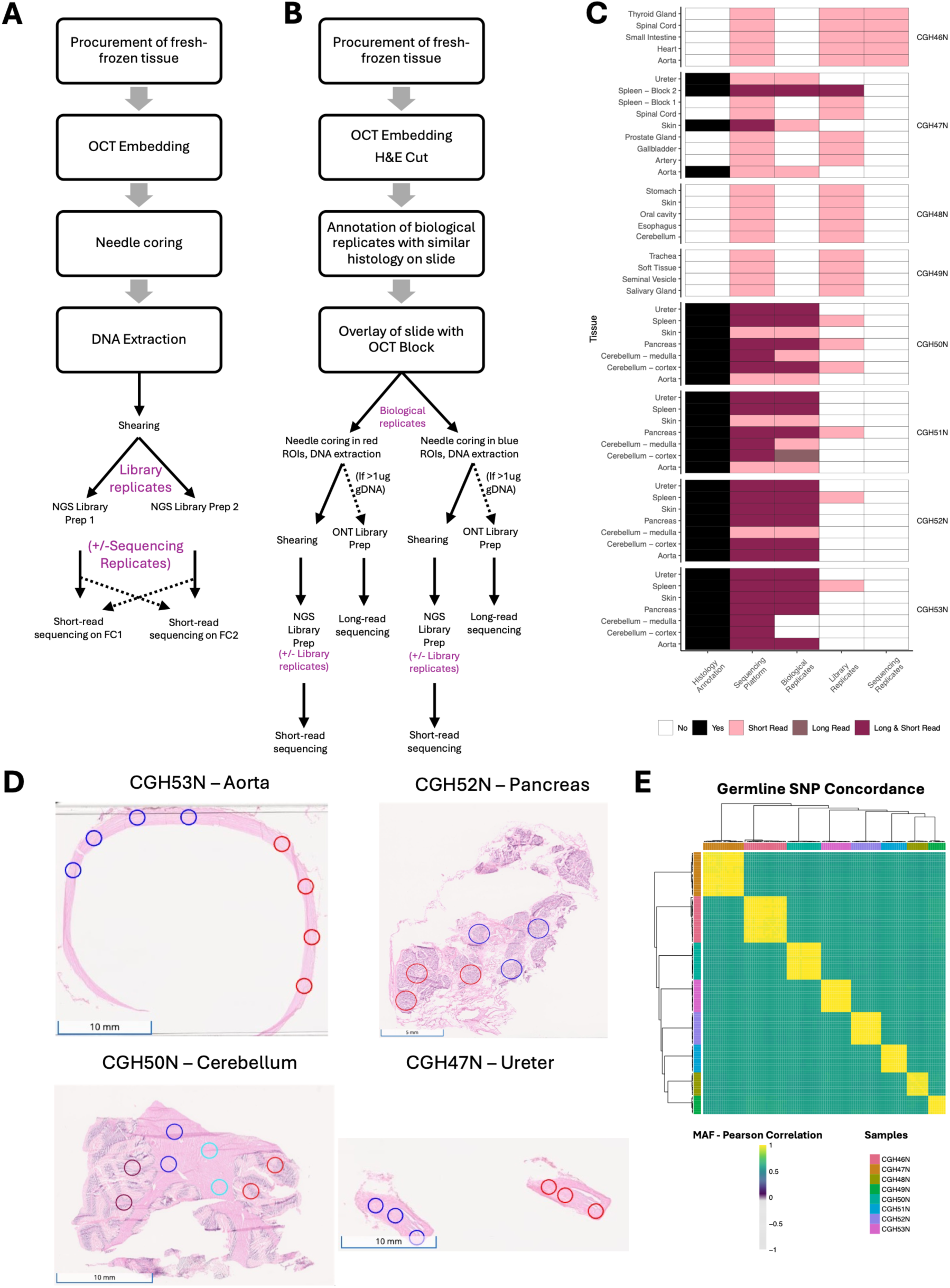
Histology, microdissection, processing and sequencing library preparation for multi-tissue samples in the JHU Cohort, related to Figure 1. **(A)** Processing and library replicate structure for tissues processed via bulk DNA extraction from needle cores without histology. **(B)** Histology and processing of tissues with macrodissected biological replicates for which DNA extraction and sequencing were conducted separately. **(C)** Tissue samples and replicates for long-read and short-read whole genome sequencing. **(D)** Examples of annotated histology slides used for microdissection of biological replicates within each tissue. **(E)** Germline SNP concordance for tissue samples from each individual illustrates expected identity for all tissues.

**Figure S2.**
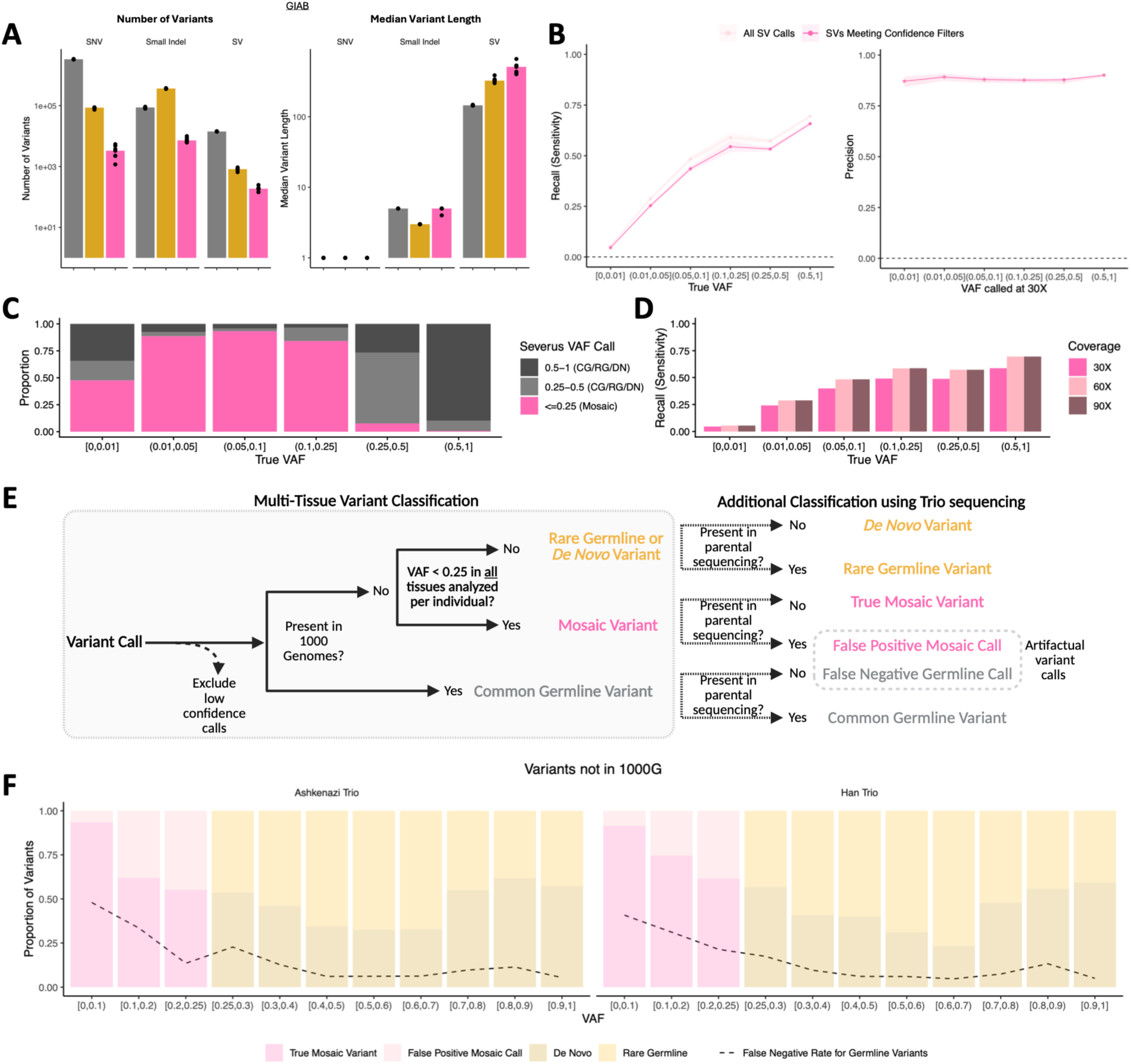
Benchmarking of long-read SV calling pipeline using Genome in a Bottle parent-offspring trios and SMaHT-MIMS benchmarks, related to Figure 2. **(A)** For Genome in a Bottle long-read sequencing data, the number (left) and median length (right) of SNVs, small indels (<50bp) and SVs (>=50bp) illustrates the biological significance of mosaic SVs. **(B)** Recall (sensitivity) and precision of our SV classification pipeline in a synthetic long-read benchmark for mosaic variant calling. The light pink line shows metrics computed using all variant calls, while the darker pink line is computed using only SVs meeting confidence filters. As expected, recall increases with VAF. **(C)** Proportion of SVs classified by our approach as Mosaic (VAF ≤ 0.25) or Germline/De Novo (VAF > 0.25) stratified by true VAF in the benchmark. The minimal number of SVs with a true VAF > 0.25 that are misclassified as mosaic highlights the accuracy of our SV classification approach. **(D)** Recall (sensitivity) of our SV calling approach demonstrates only modest improvements with increased sequencing coverage. **(E)** Schematic illustrating how parent-offspring trio data can be used to disambiguate between *de novo* and rare germline variants, and to quantify rates of false positive mosaic calls (appear subclonal in the offspring despite confirmation in parental sequencing) and false negative germline calls (missed in the parent despite being a common germline variant). **(F)** Proportions of true mosaic vs. false positive mosaic SV calls, and *de novo* vs. rare germline SVs as disambiguated using two parent-offspring trios within the GIAB cohort. In all cases, the proportions of true mosaic SVs and *de novo* SVs exceed the false negative germline rate suggesting that *bona fide* somatic SVs occur frequently.

**Figure S3.**
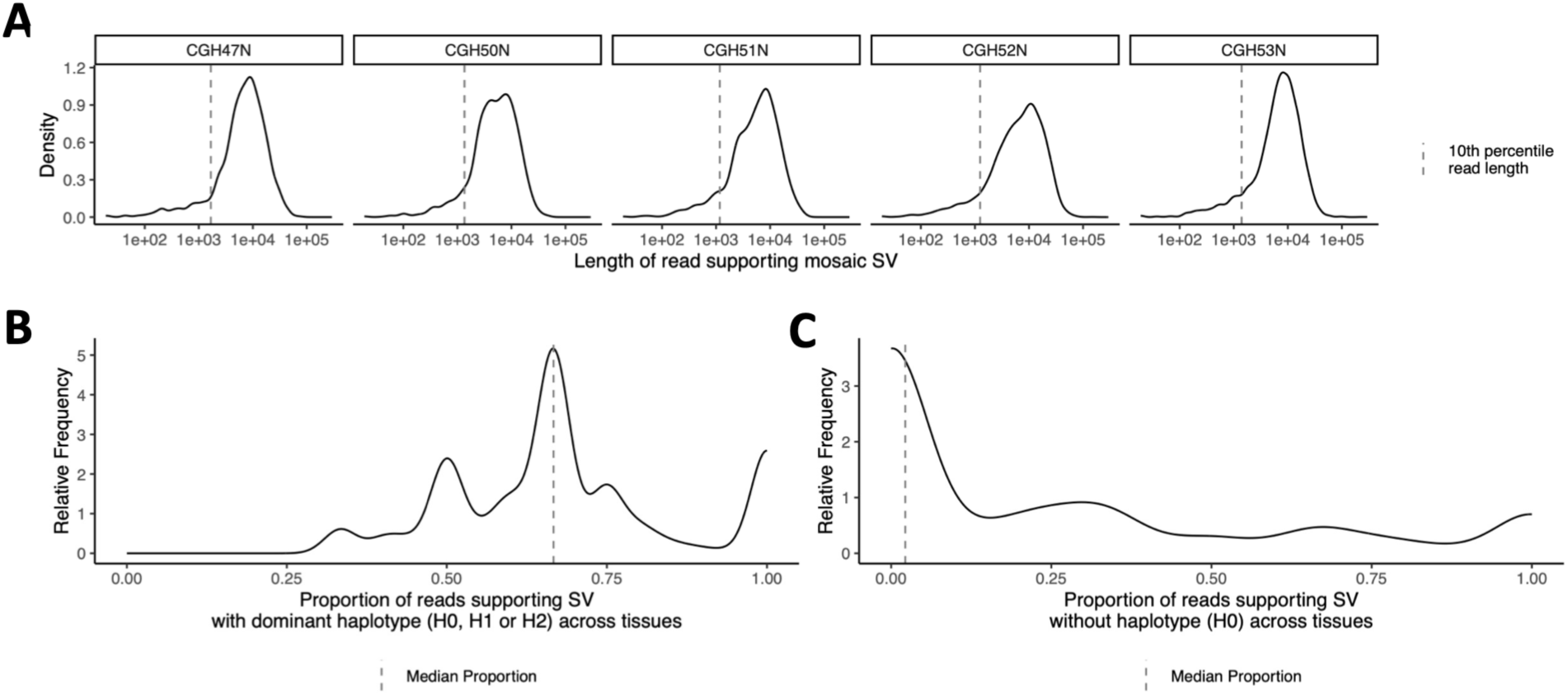
Read-level metrics for reads tagged as supporting a mosaic SV within 1000bp upstream or downstream of the breakpoint, related to Figure 2. **(A)** Distribution of supporting read lengths for all putative mosaic SVs in the JHU Cohort (n=2610). **(B)** For all putative mosaic variants, the distribution of proportion of reads supporting the SV with the dominant haplotype across tissues (defined as the haplotype assigned in the largest number of tissues for that variant) **(C)** For all putative mosaic variants, the proportion of reads not haplotyped.

**Figure S4.**
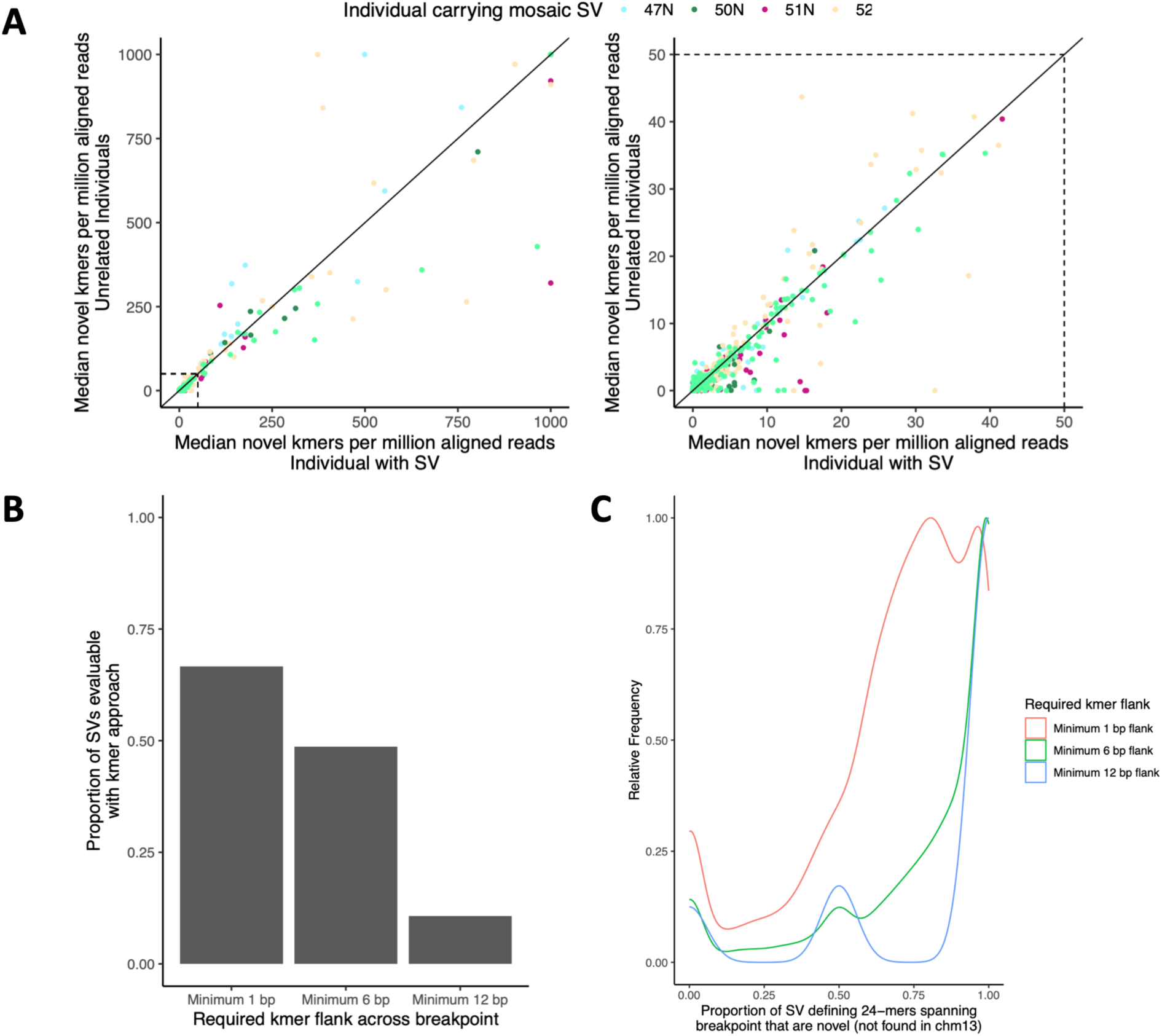
Kmer-based short-read sequencing approach for orthogonal validation of mosaic SVs discovered with long-read sequencing, related to Figure 2. **(A)** Median counts of novel SV breakpoint kmers in samples from the individual with the SV (x-axis) and unrelated individuals (y-axis) highlight the enrichment of these kmers in individuals carrying the mosaic SV. Novel kmers are those not found in the chm13 reference genome and formed by an SV breakpoint. Points below the diagonal indicate validated SVs (kmer count increased in the individual with the SV). Each point represents kmer counts for a single mosaic SV found in a single individual. The right panel is a zoomed inset of the left panel in the range 0 to 50 novel kmers per million aligned reads (indicated by the dotted lines). **(B)** The proportion of evaluable mosaic SVs decreases as the minimum required flank for novel kmers on either side of the SV breakpoint increases. An SV is considered evaluable if atleast 1 novel breakpoint kmer is found in atleast 1 sample. A novel kmer is one not found in the chm13 reference genome and formed by a SV breakpoint. At a minimum 1bp, any novel kmer spanning the breakpoint is counted. For a minimum 6bp flank, atleast 6bp must be found on either side of the breakpoint. For a minimum 12bp flank, 24bp kmers must be centered across the breakpoint. **(C)** Distribution of the proportion of kmers spanning mosaic SV breakpoints that are novel (not found in chm13), stratified by minimum required flank on either side of the breakpoint. Most kmers spanning mosaic SV breakpoints are novel, but a substantial fraction exists that are found elsewhere in the reference genome consistent with the non-unique nature of short kmers.

**Figure S5.**
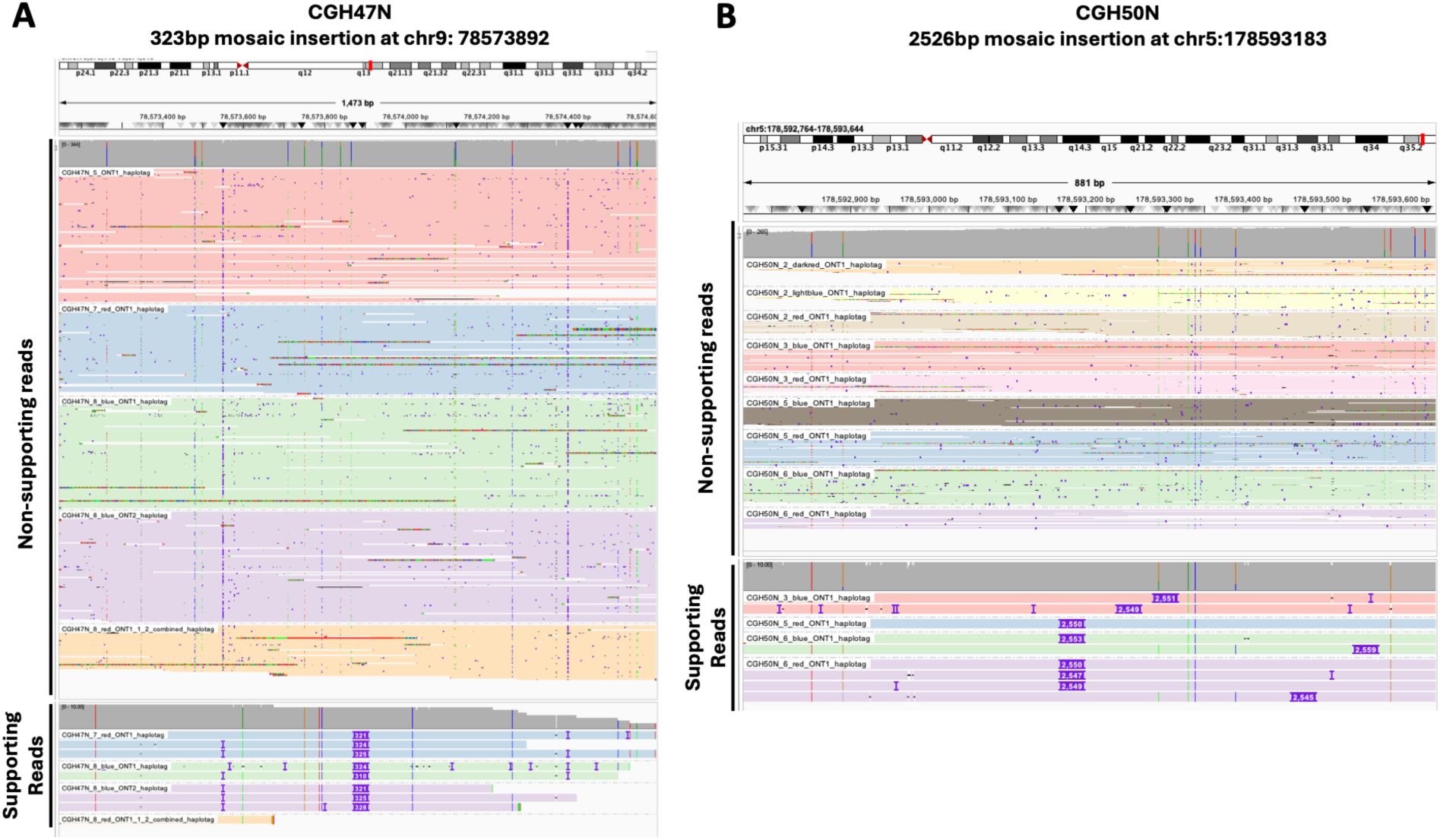
Examples of mosaic insertions in the JHU Cohort, related to Figures 2 and 4. **(A)** IGV screenshot showing example of mosaic insertion in CGH47N. Mechanism of repeat element involvement for this SV is shown in Figures 4C and 4D. **(B)** IGV screenshot showing example of mosaic insertion in CGH50N. Mechanism of repeat element involvement for this SV is shown in Figures 4E and 4F. IGV screenshots of non-supporting reads spanning the breakpoint are shown in a collapsed view, grouped and colored by tissue sample of origin. Reads supporting the SV are shown in an expanded view, with the same colors for each tissue sample and the insertion marked. In (B) some imprecision in breakpoint location is noted due to repetitive alignment. Supporting and non-supporting reads are categorized as such by the variant caller.

**Figure S6.**
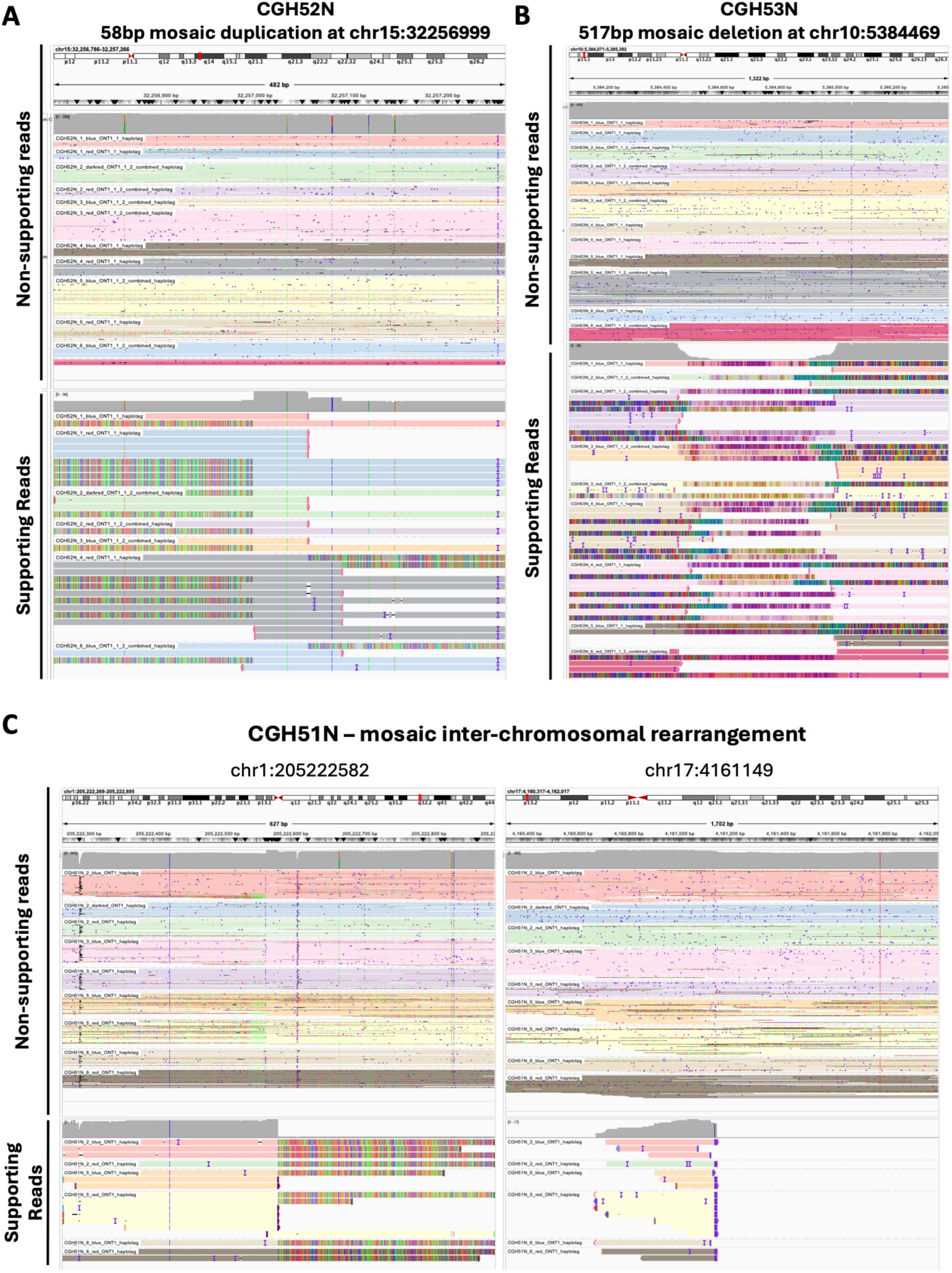
Examples of mosaic SVs in the JHU Cohort, related to Figure 2. IGV screenshots of example mosaic SVs: **(A)** duplication, **(B)** deletion, and **(C)** inter-chromosomal rearrangement. Non-supporting reads spanning the breakpoint are shown in a collapsed view, grouped and colored by tissue sample of origin. Reads supporting the SV are shown in an expanded view, with the same colors for each tissue sample and soft- and hard- clipped support indicated. Supporting and non-supporting reads are categorized as such by the variant caller.

**Figure S7.**
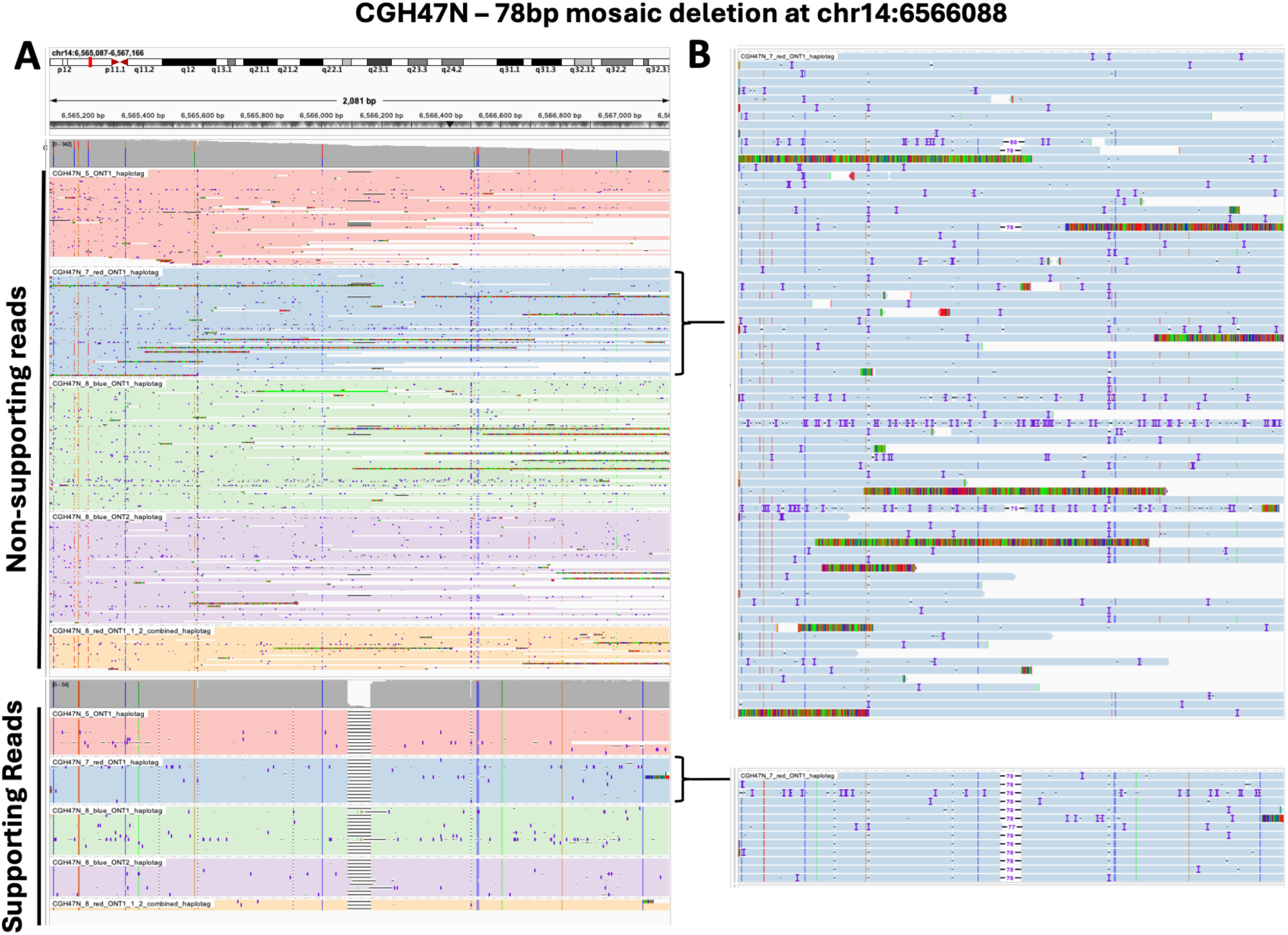
Example of a mosaic deletion in the JHU Cohort, related to Figure 2. **(A)** IGV screenshots of non-supporting reads spanning the breakpoint are shown in a collapsed view, grouped and colored by tissue sample of origin. Reads supporting the SV are shown below, also in a collapsed view, with the same colors for each tissue sample. **(B)** Expanded view of non-supporting and supporting reads from one tissue sample highlighting that the mosaic SV may also occur in a few reads not tagged as supporting by the SV caller due to quality or length, but that the frequency of these reads is rare and does not exceed a mosaic VAF. Supporting and non-supporting reads are categorized as such by the variant caller.

**Figure S8.**
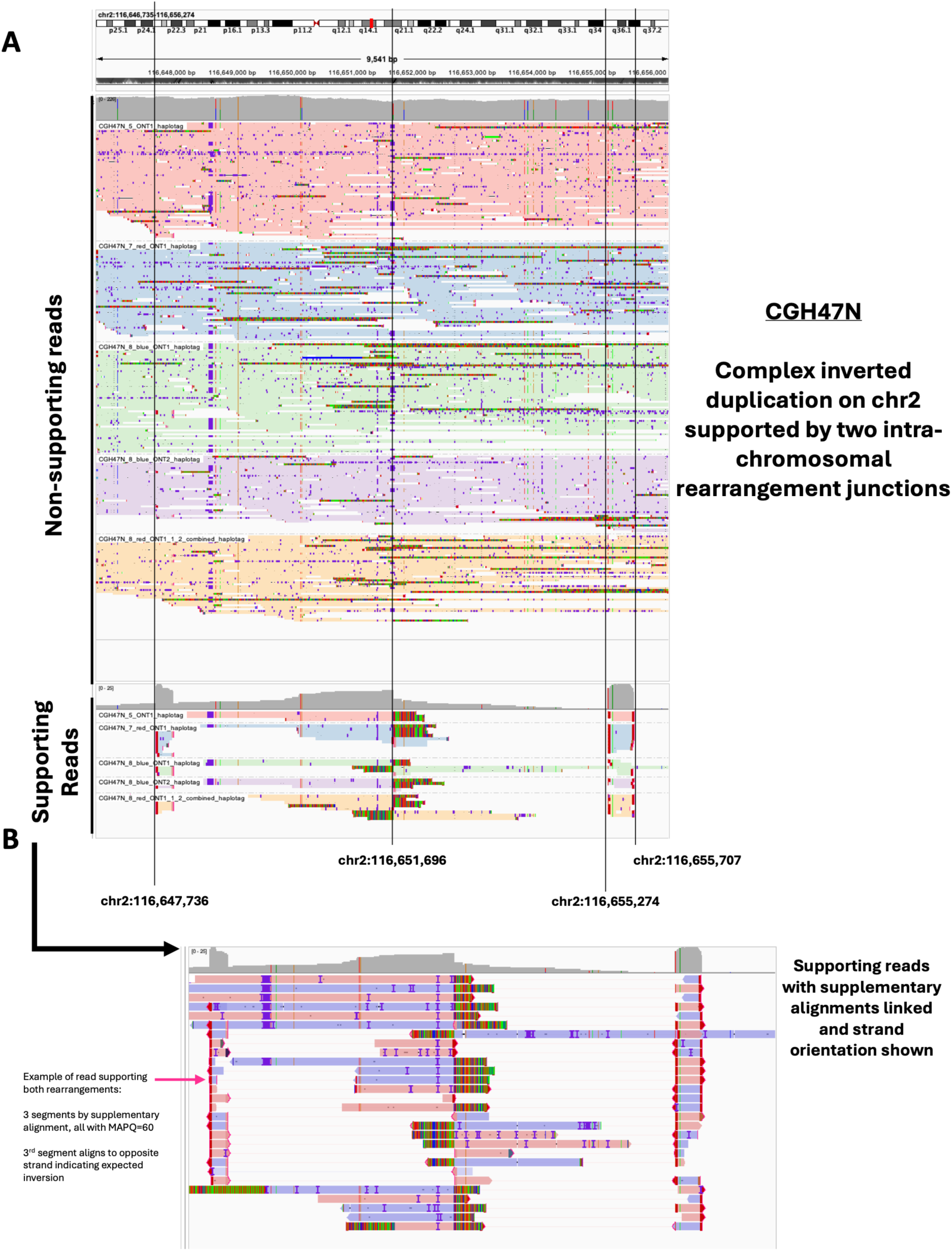
Example of a complex cluster of mosaic intra-chromosomal rearrangements in the JHU Cohort, related to Figure 2. **(A)** IGV screenshots of non-supporting reads spanning 4 breakpoints are shown in a collapsed view, grouped and colored by tissue sample of origin. Reads supporting the SV are shown below, also in a collapsed view, with the same colors for each tissue sample. **(B)** Expanded view of supporting reads colored by strand orientation of alignment, with supplementary alignments linked, shows an inversion duplication. Supporting and non-supporting reads are categorized as such by the variant caller.

**Figure S9.**
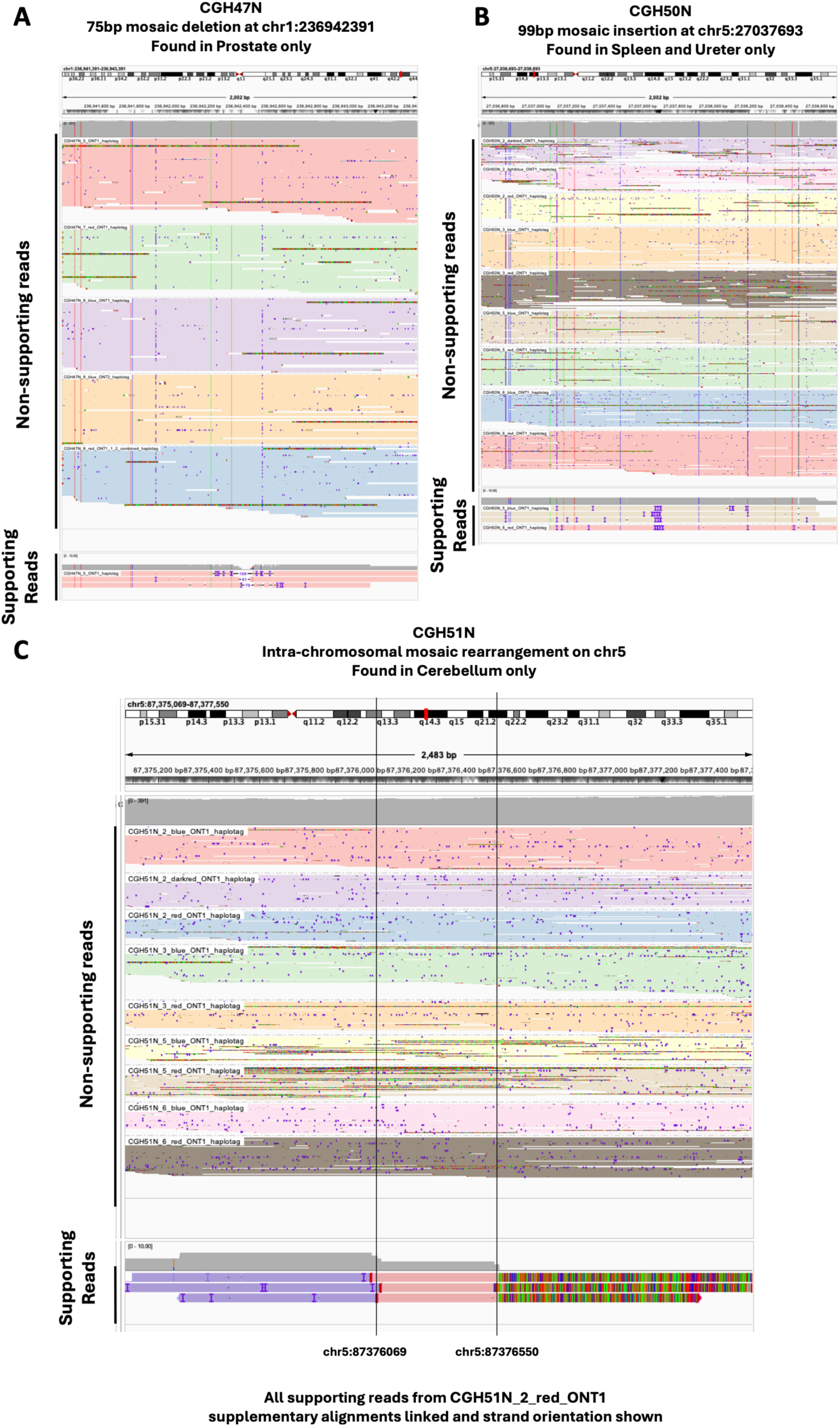
Examples of tissue-restricted mosaic SVs in the JHU Cohort, related to Figure 2. IGV screenshots of example mosaic SVs: **(A)** Deletion found in prostate only, **(B)** Insertion found in spleen and ureter only, and **(C)** intra-chromosomal rearrangement found in cerebellum only. Non-supporting reads spanning the breakpoint are shown in a collapsed view, grouped and colored by tissue sample of origin. Supporting and non-supporting reads are categorized as such by the variant caller. For (A) and (B), reads supporting the SV are shown in an expanded view, with the same colors for each tissue sample and soft- and hard- clipped support indicated. For (C), supporting reads are colored by strand orientation of alignment, with supplementary alignments linked, showing the expected reversal in strand orientation.

**Figure S10.**
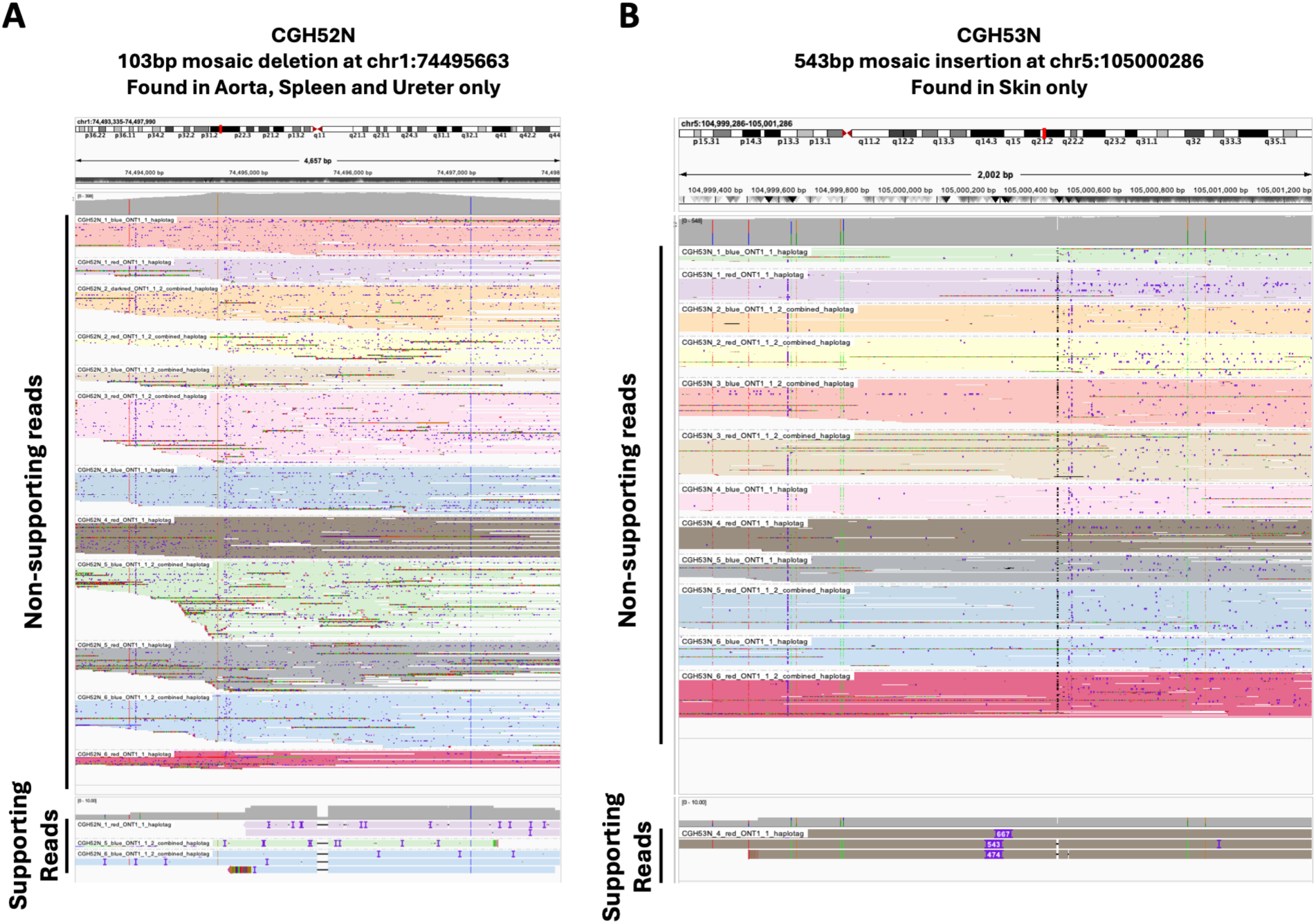
Additional examples of tissue-restricted mosaic SVs in the JHU Cohort, related to Figure 2. IGV screenshots of example mosaic SVs: **(A)** Deletion found in aorta, spleen and ureter only and **(B)** Insertion found in skin only. Non-supporting reads spanning the breakpoint are shown in a collapsed view, grouped and colored by tissue sample of origin. Supporting and non-supporting reads are categorized as such by the variant caller. Reads supporting the SV are shown in an expanded view, with the same colors for each tissue sample and soft- and hard- clipped support indicated.

**Figure S11.**
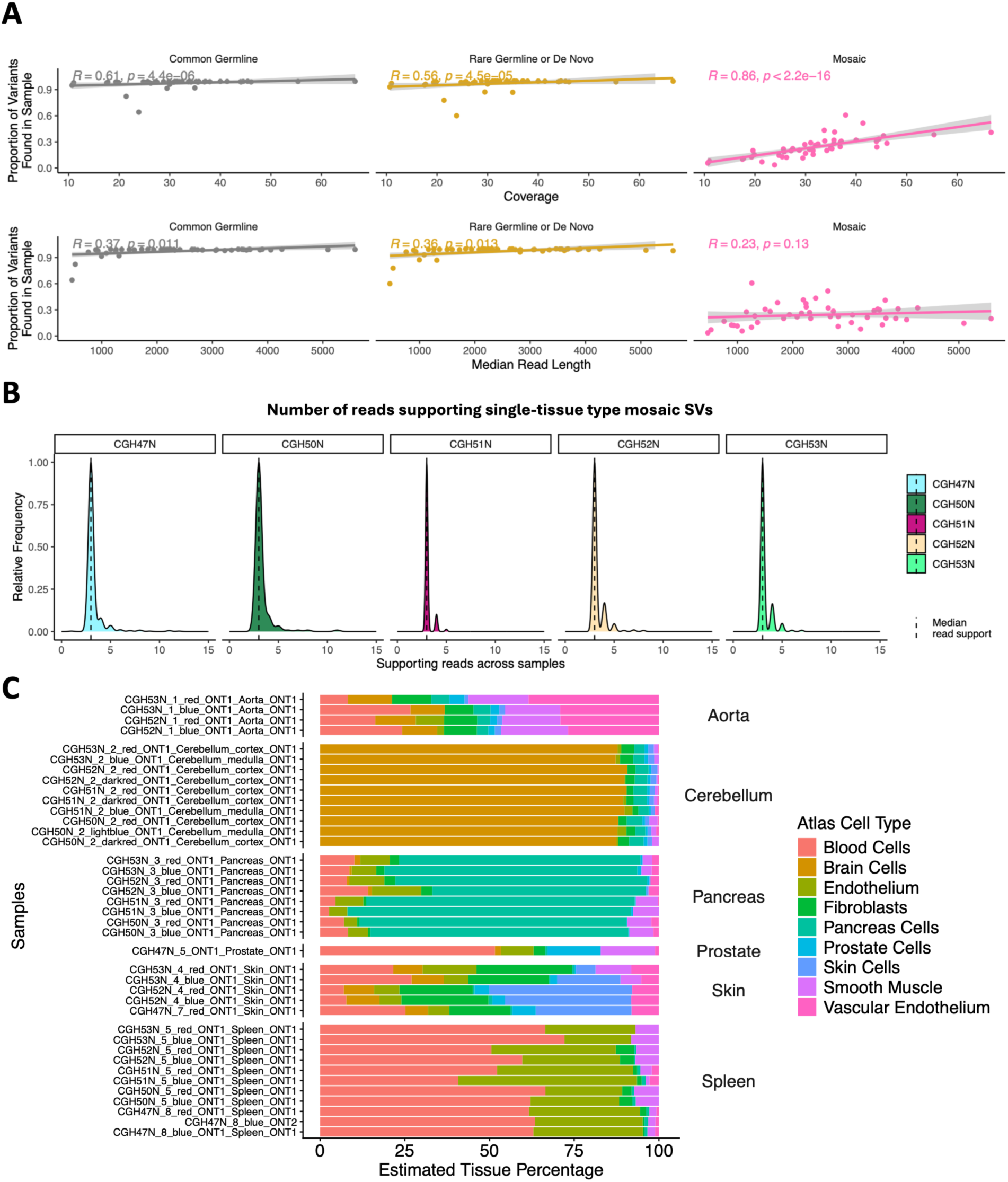
Characterization of SV call read and tissue support in the JHU Cohort, related to Figure 3. **(A)** Proportion of variants in the call set found in each tissue sample (vertical axis) as a function of sequencing coverage and median read length (horizontal axis) suggests that technical missingness may have some influence on patterns of tissue-restricted variants. **(B)** Distribution of number of long reads supporting single tissue mosaic SVs. Mosaic SVs found in only a single tissue type are supported by a median of 3 long reads. **(C)** Methylation deconvolution of cell-type contributions to tissue samples used for long-read mosaic SV identification. The majority of samples exhibit tissue-type specific identity, while spleen samples have high blood cell contribution likely due to expected immune infiltration. Ureter samples are not shown as no concordant cell types were included in the published methylation atlas^82^ utilized for this analysis.

**Figure S12.**
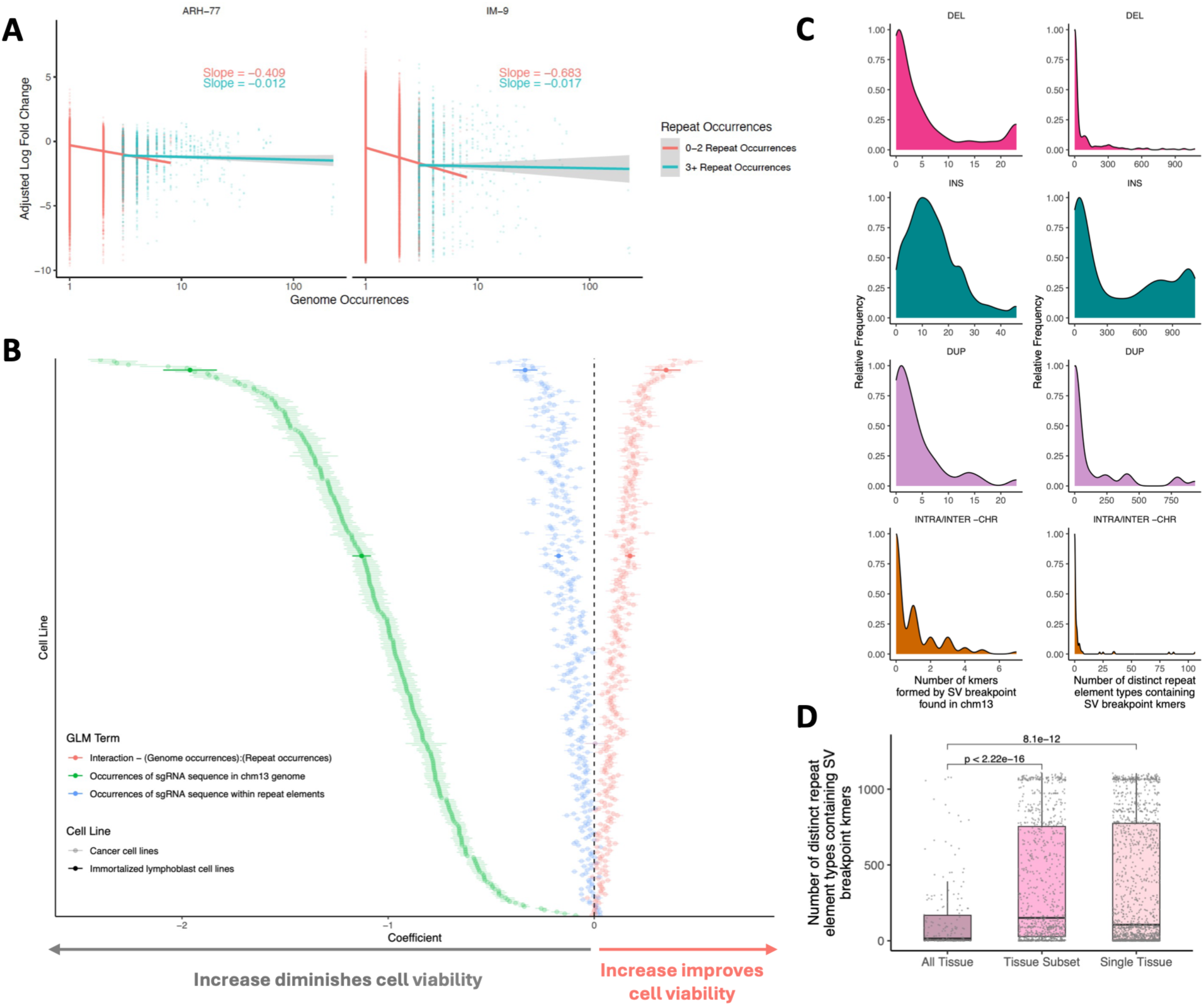
Deleterious impacts of double-stranded breaks in the genome are attenuated when occurring within repeat elements, related to Figure 4. **(A)** Relationship between number of genome occurrences (DSB cut-sites) and adjusted log-fold-change for each sgRNA in a CRISPR essentiality screen of 2 immortalized cell lines. Negative log-fold-change indicates a more deleterious impact, and the flattening of the fitted slope for sgRNA sequences with more occurrences within annotated repeat elements suggests a less deleterious impact. **(B)** Coefficients for fitted model in all screened cell lines (n=318) illustrate that higher numbers of introduced DSBs have more negative impacts (negative coefficients) on cell viability but that this impact is attenuated by occurrences within annotated repeat elements (positive coefficients for interaction term). **(C)** Distribution of the number of kmers formed by mosaic SV breakpoints that are found within the chm13 reference genome (left), and the number of repeat element types that they map to (right). Many mosaic SV breakpoints form short 24bp sequences (kmers) with sequence homology corresponding to an annotated repeat element type. **(D)** The number of repeat element types containing an SV breakpoint kmer is higher for mosaic SVs found in single tissues or subsets of tissues as compared to early embryonic mosaic SVs found in all tissues of an individual. Each point represents a mosaic SV.

**Figure S13.**
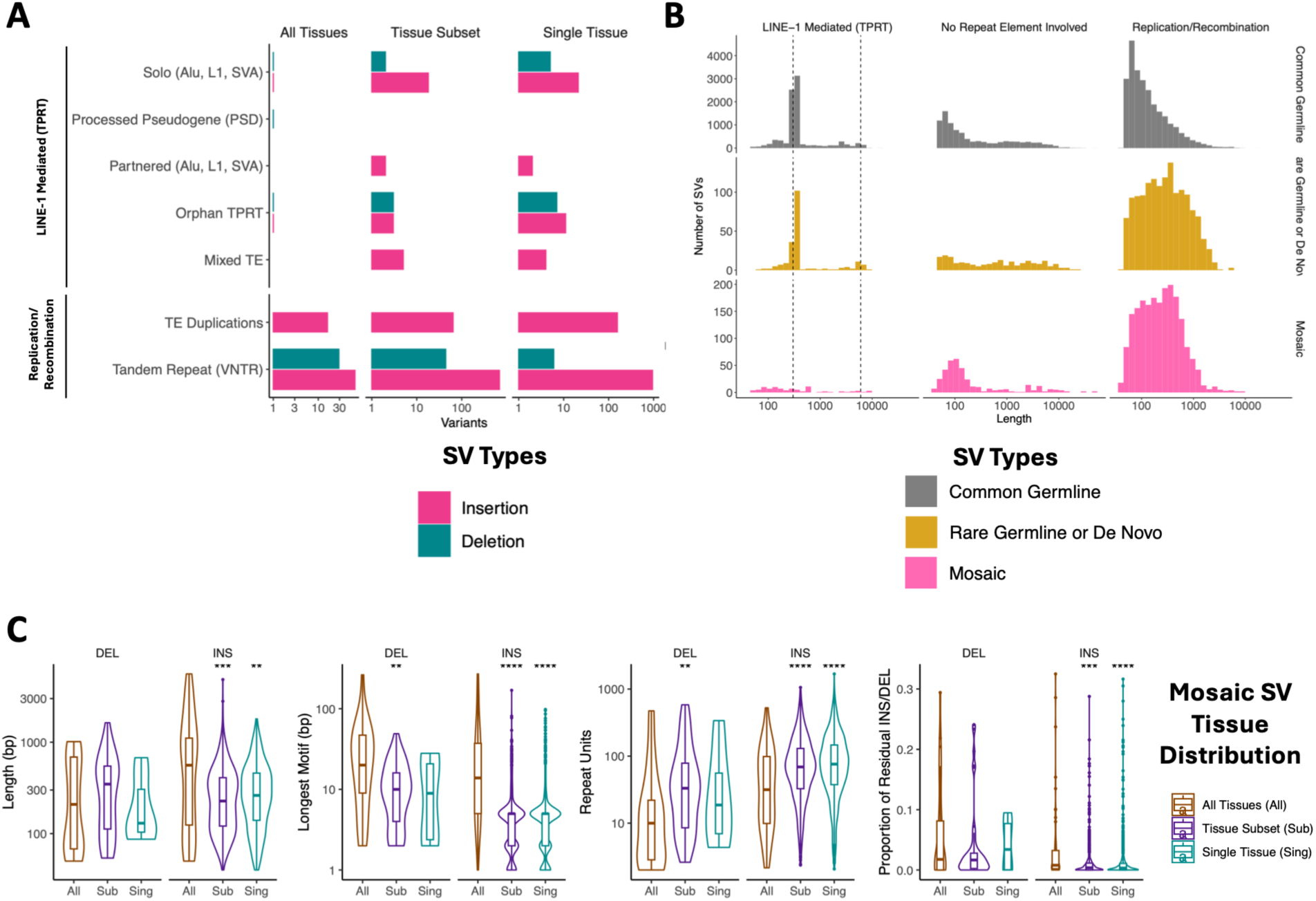
Additional characterization of repeat element mediation of mosaic SV calls in the JHU Cohort, related to Figure 4. **(A)** Detailed annotation of mechanism for repeat-element involved mosaic insertion and deletion SV calls in the JHU Cohort, stratified by observed tissue distribution. **(B)** Length distributions for insertion and deletion SV calls in the JHU cohort stratified by signature of repeat element involvement (LINE-1 Mediated or replication/recombination) and type (Common Germline, Rare Germline/*de novo*, Mosaic). **(C)** Characteristics of mosaic variable number tandem repeat (VNTR) insertions and deletions in the JHU Cohort, stratified by observed tissue distribution. The total SV length (panel 1, leftmost), longest dominant motif (panel 2), number of repeat units (panel 3) and proportion of residual incomplete sequence (panel 4, rightmost) varies by tissue distribution of mosaic SVs, with the longest lengths and highest number of repeat units generally found in mosaic SVs restricted to single tissues or subsets of tissues.

**Figure S14.**
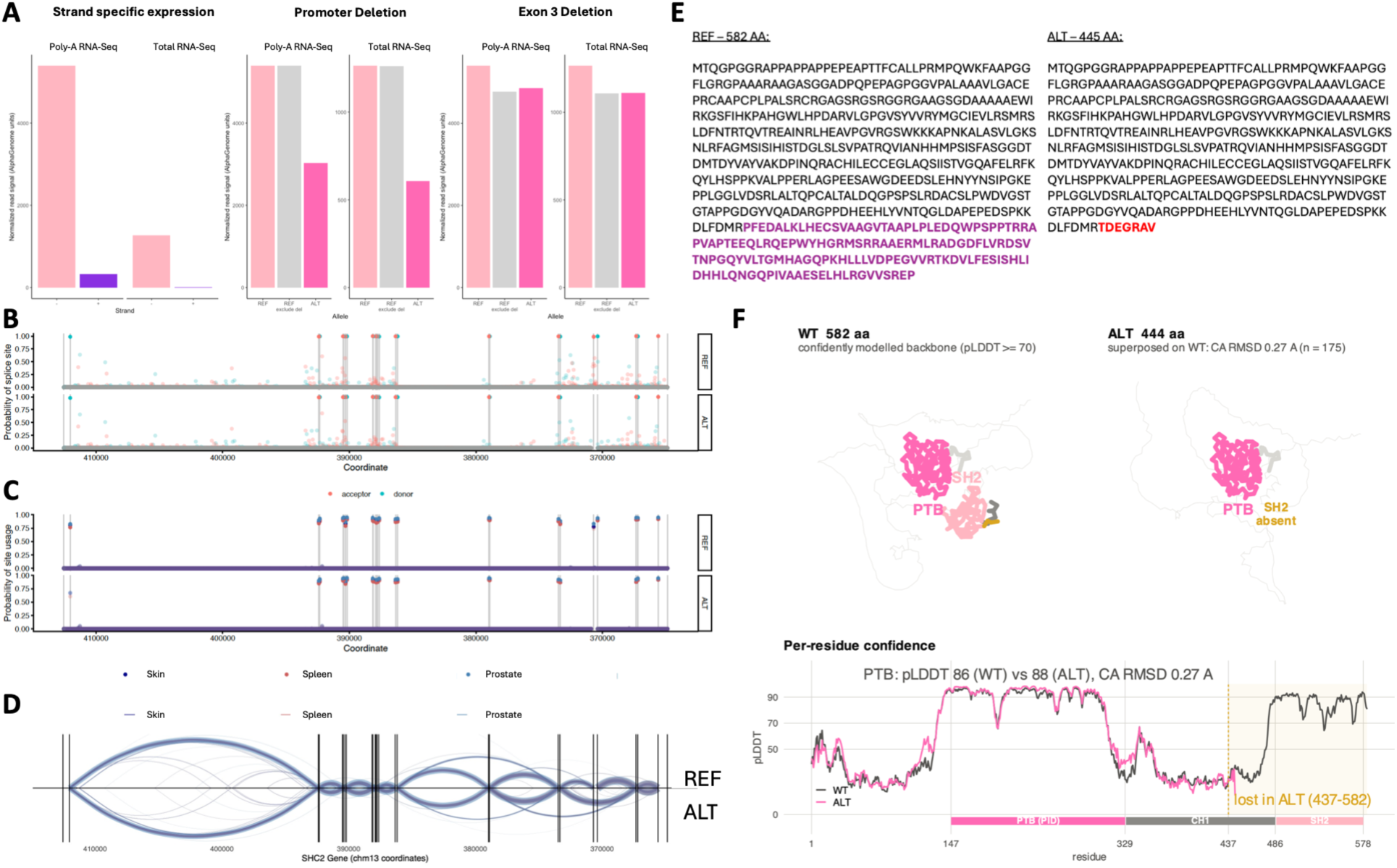
AlphaGenome predictions reveal putative functional consequences for mosaic deletion of SHC2 exon 3, related to Figure 5. **(A)** Predicted SHC2 gene expression for WT and variant alleles demonstrates biological plausibility of AlphaGenome predictions in this genomic region. Left panel, predicted SHC2 expression shows the expected negative strand specificity. Middle and right panels, predicted loss of expression from SHC2 promoter deletion (middle) and exon 3 deletion (right). The light pink bar shows predicted gene expression for the REF allele. The gray bar shows predicted expression for the REF sequence with the signal contributed from deleted bases subtracted. The dark pink bar shows predicted expression for the ALT sequence containing the deletion. As expected, a promoter deletion greatly diminishes gene expression signal despite the minimal signal contribution of this region while an exon 3 deletion shows a reduction in expression proportional to the amount of deleted sequence. Predictions are shown for lung tissue, which natively expresses SHC2. **(B)** Predicted presence of donor and acceptor splice sites across the SHC2 gene body in the REF and ALT alleles align with known exon boundaries and show the expected splice site loss surrounding the mosaic exon 3 deletion in the ALT allele. Predictions are shown for skin, spleen and prostate – the three tissues sequenced for the individual, CGH47N, with this mosaic SV. **(C)** Predicted usage of splice sites across the SHC2 gene body in the REF and ALT alleles confirms that the exon 3 boundary splice sites are used in transcription of the REF transcript, but lost in the ALT. **(D)** Sashimi plot showing predicted splice junctions across the SHC2 gene body in the REF and ALT alleles shows a novel splice junction between exon 2 and exon 4 created by deletion of exon 3. Arc thickness is proportional to the predicted junction usage score. **(E)** Amino acid sequences encoded by the REF and ALT transcripts, showing how mosaic exon 3 deletion leads to a premature stop codon and truncation. **(F)** AlphaFold predictions for the amino acid sequences in (E) show the nonfunctional protein predicted to result from the mosaic SHC2 deletion observed in the JHU cohort. This protein is missing the SH2 domain and is truncated sufficiently that it likely triggers nonsense mediated decay.

**Figure S15.**
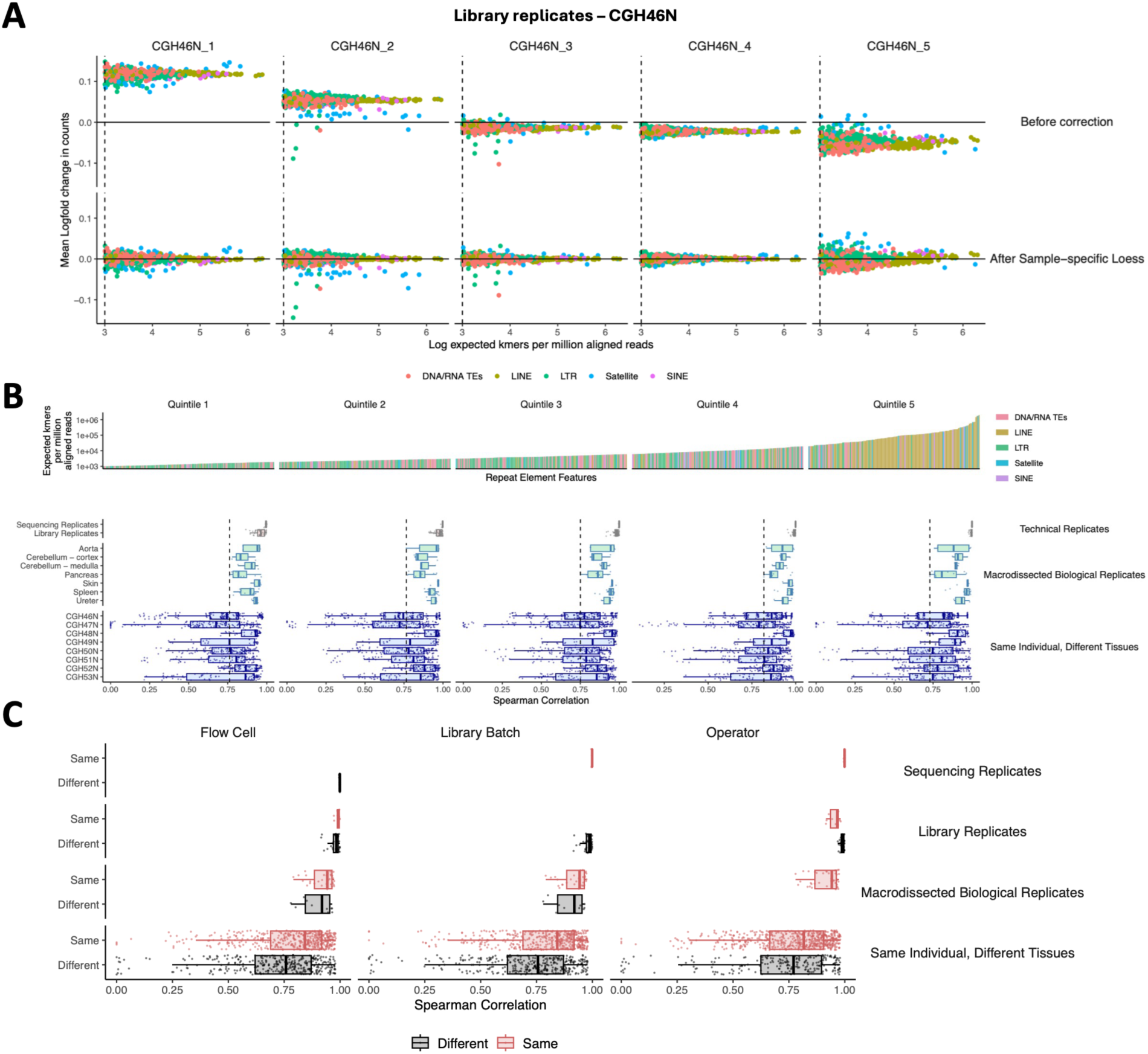
Characterization of intra-individual variation in genome-wide repeat landscapes constructed from short-read sequencing, related to Figure 6. **(A)** Examples of kmer repeat landscape comparisons between library replicates before (top row) and after (bottom row) application of a sample-specific loess correction. For each alignment-free kmer repeat landscape, technical variation in element counts due to sequencing coverage and duplicates is regressed out prior to comparison. **(B)** Increased variation in repeat landscapes between different tissues from the same individual is observed across all tissues and individuals studied, as well as for all quintiles of repeat element counts. Each point represents the Spearman’s correlation coefficient for a given comparison pair and repeat element quintile. Kmer repeat landscapes comprise 786 individual repeat elements, divided into five quintiles by expected abundance in the genome. **(C)** Repeat landscapes exhibit high concordance in replicates and patterns of variation across different tissues regardless of technical factors including sequencing flow cell, library batch and operator. Each point represents the median Spearman’s correlation coefficient across repeat element quintiles for a given comparison pair.

**Figure S16.**
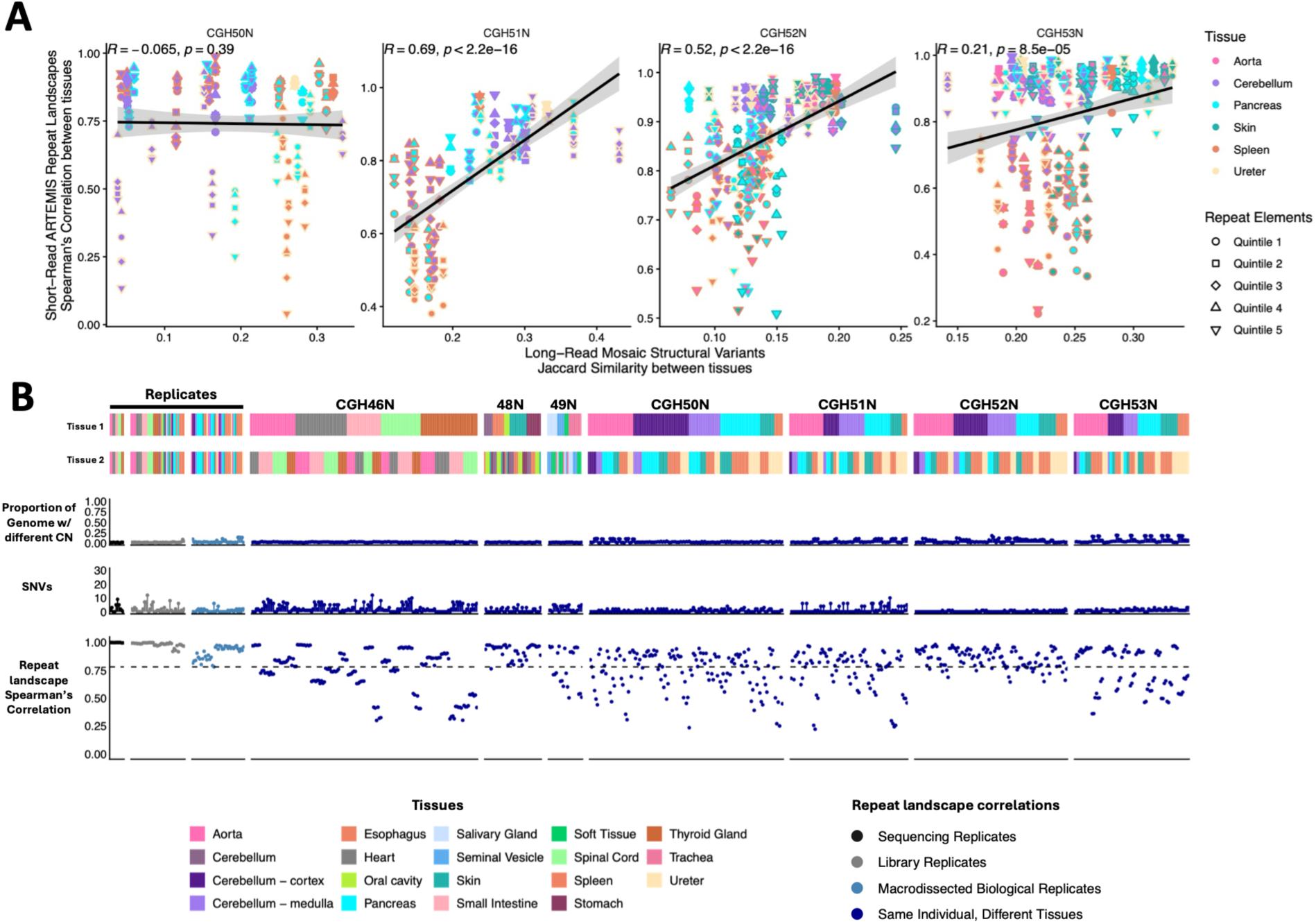
Comparisons of global repeat landscapes reveal widespread tissue-specific mosaicism across normal tissues, related to Figure 6. For individuals CGH50N, 51N, 52N and 53N (47N is shown in Figure 6): **(A)** Tissue similarity as computed from short-read ARTEMIS repeat landscapes is broadly concordant with similarity computed from long-read mosaic SVs. For each pair of tissues, the y-axis shows repeat landscape correlation, and the x-axis shows Jaccard similarity between mosaic SV calls. The individual without concordance (CGH50N) is also the individual with the least number of mosaic SVs identified. **(B)** For each pairwise tissue sample comparison, copy number differences, SNVs and repeat landscape correlations reveal tissue-specific repeat element differences to be the main driver of variation.

**Figure S17.**
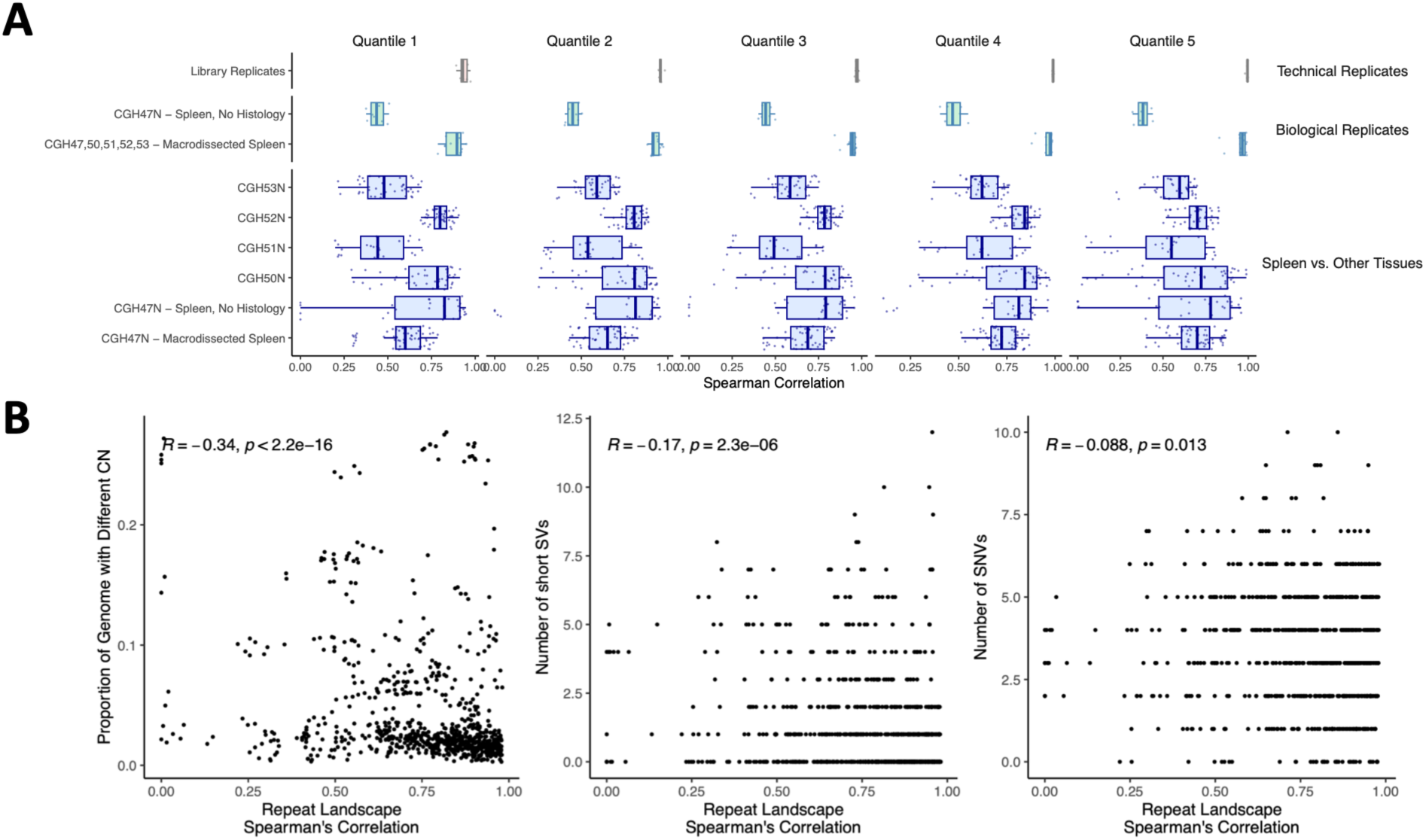
Impact of biological sources of variation on intra-individual repeat landscape concordance, related to Figure 6. **(A)** For individual CGH47N, macrodissected spleen tissue replicates and bulk DNA extract replicates without histology are available. Macrodissected replicates appear more similar to each other than bulk replicates as expected; however in cross-tissue comparisons all spleen samples across individuals exhibit high patterns of variation as compared to other tissue types. **(B)** Relationship between repeat landscape concordance (horizontal axis) and copy number alterations (left panel, vertical axis), short SVs not within repeat elements that are able to be called in short-read sequencing (vertical axis, middle panel), and SNVs called from short-read sequencing (right panel, vertical axis) illustrate that tissue-specific variation in repeat landscapes is widespread including in tissues considered similar by other metrics. Each point represents a pairwise comparison between different tissues from an individual.

**Figure S18.**
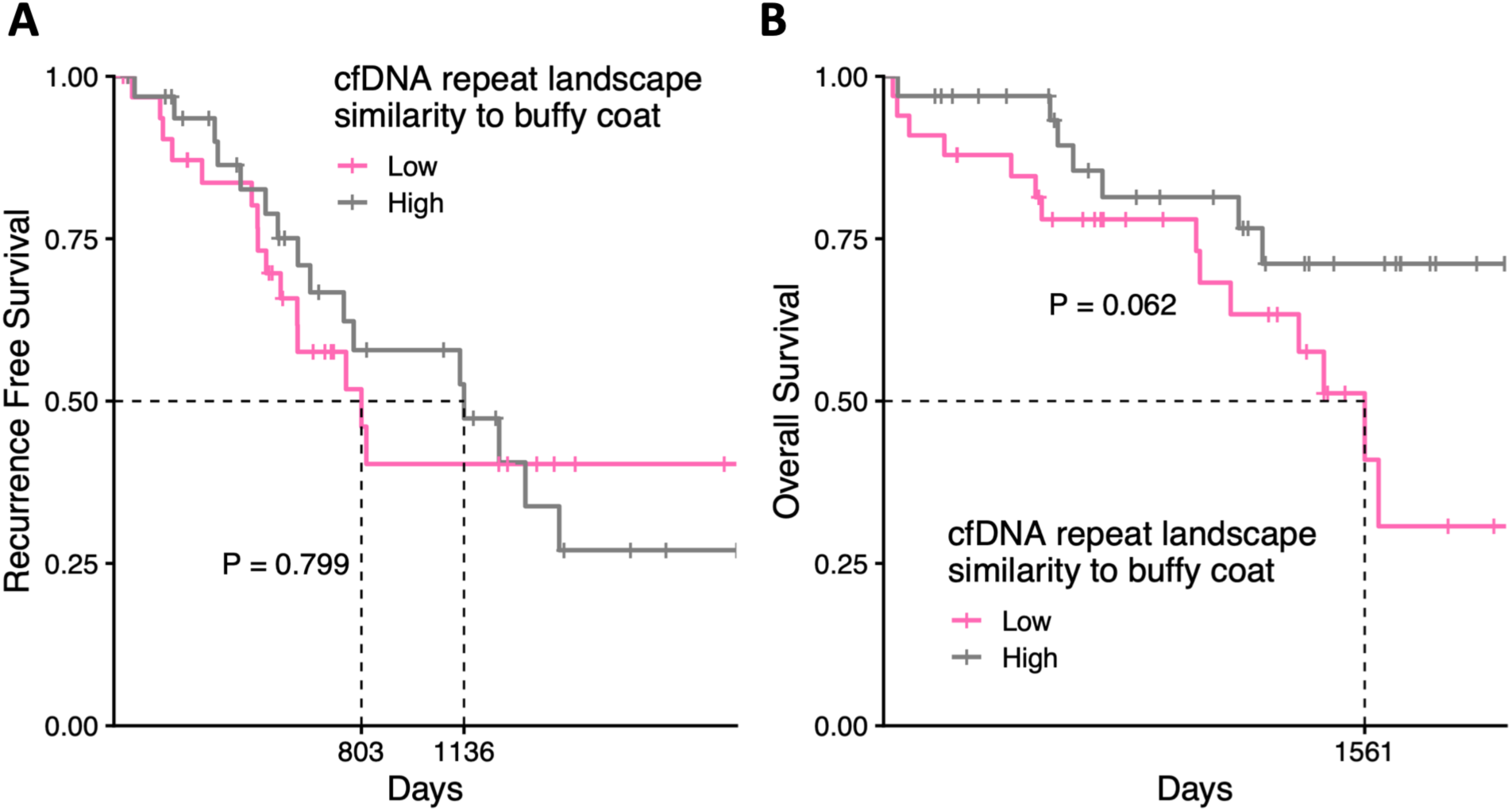
At the baseline pre-treatment timepoint, repeat landscape correlations between cfDNA and buffy coat in patients with ovarian cancer do not correlate with survival, related to Figure 7. **(A)** Kaplan-Meier curves for recurrence free survival in patients with ovarian cancer based on baseline pre-treatment repeat landscape correlations between plasma and buffy coat. **(B)** Similar to (A), but for overall survival.

